# Divergence time and environmental similarity predict the strength of morphological convergence in stick and leaf insects

**DOI:** 10.1101/2023.11.07.565940

**Authors:** Romain P. Boisseau, Sven Bradler, Douglas J. Emlen

## Abstract

Independent evolution of similar traits in lineages inhabiting similar environments (convergent evolution) is often taken as evidence for adaptation by natural selection, and used to illustrate the predictability of evolution. Yet convergence is rarely perfect. Environments may not be as similar as they appear (e.g., habitats scored the same may be heterogenous to the organisms). And lineages can evolve in different ways even when submitted to the same environmental challenges, because responses to selection are contingent upon available genetic variation and independent lineages may differ in the alleles, genetic backgrounds, and even the developmental mechanisms responsible for the phenotypes in question. Both impediments to convergence are predicted to increase as the length of time separating two lineages increases, making it difficult to discern their relative importance. We quantified environmental similarity and the extent of convergence to show how habitat and divergence time each contribute to observed patterns of morphological evolution in stick and leaf insects (order Phasmatodea). Dozens of phasmid lineages independently colonized similar habitats, repeatedly evolving in parallel directions on a 26-trait morphospace, though the magnitude and direction of these shifts varied. Lineages converging towards more similar environments ended up closer on the morphospace, as did closely related lineages, and closely related lineages followed more parallel trajectories to arrive there. Remarkably, after accounting for habitat similarity, we show that divergence time reduced convergence at a constant rate across more than 60 million years of separation, suggesting even the magnitude of contingency can be predictable, given sufficient spans of time.

**Significance statement:** Phasmids (stick and leaf insects) exemplify the extraordinary power of natural selection to shape organismal phenotypes. The animals themselves are charismatic champions of crypsis and masquerade; and our characterization of their adaptive radiation reveals dozens of instances of convergence, as lineages adapted to similar changes in habitat by repeatedly evolving similar body forms. Our findings show that the similarity of environmental conditions experienced by the organisms – the closeness of the invaded niches – and the extent of elapsed time since divergence, both predict the strength of morphological convergence. The phasmid radiation reveals an evolutionary process that is surprisingly predictable, even when lineages have been evolving independently for tens of millions of years.

## Introduction

When does convergent evolution happen? Examples of lineages independently evolving similar phenotypes are numerous and conspicuous (1–4) (e.g., gliding mammals (5), cave amphipods (6, 7), Hawaiian spiders (8)), and likely result from adaptation to similar ecological niches (9, 10; but see (11)). Yet convergence is rarely perfect and sometimes does not occur at all, even when habitats are similar. When it does occur, the extent of phenotypic similarity varies widely (5, 9, 12) and the factors causing this variation and, by extension, influencing the repeatability of evolutionary outcomes, are not well understood (13).

One important determinant of the likelihood and extent of convergence is the degree of relatedness among lineages. Repeated evolution usually involves closely related taxa (10)(e.g., Caribbean *Anolis* lizards (14, 15), three-spined stickleback fish (16)), suggesting that strong convergence is most likely when the time separating lineages is brief. Gould famously argued that evolutionary outcomes are contingent on the intricate series of historical events uniquely experienced by each lineage (17–19). Closely related lineages share more of their evolutionary history and, consequently, more of their genetic variation (16, 20–25). Closely related lineages are also more likely to share the same ancestral niche and associated ancestral phenotypes (13). Threespine stickleback repeatedly colonized lakes and streams from the same marine habitat (26, 27). In these instances, adaptation to the new niche is likely to proceed through similar sequences of phenotypic changes (i.e., parallel evolutionary trajectories) arriving at phenotypes that are strongly convergent. More disparate lineages may approach a shared environmental challenge from different starting phenotypes, with weaker convergence as a result. And lineages with enough accumulated differences may not converge at all. Aye-ayes (Primates) and woodpeckers (Aves) each catch and eat insect larvae found under the bark of trees, yet they forage in strikingly different ways (13). Aye-ayes use their teeth to break through the bark and an elongated middle finger to catch larvae, while woodpeckers use hammering beaks to get through the bark and long, barbed tongues to catch insects.

Consequently, the extent of shared evolutionary history and the similarity of phenotypic ancestral states should each affect the likelihood of strong convergence, such that the greater the opportunity for contingency – the more accumulated time since their split -- the less likely any two lineages should be to converge strongly in response to a shared selection environment. Yet rigorous tests of Gould’s prediction are lacking, in part because examples of strong convergence between distantly related lineages are rare, and also because the majority of studies of repeated evolution focus on lineages with minimal (<5 million years) time since divergence, so little opportunity for contingency.

Quantifying the role of divergence time on convergence requires a system: (i) where the extent of phenotypic convergence and environmental similarity can be quantified precisely; (ii) where instances of convergence span vast periods of time from recently diverged to much more distantly genetically related lineages; and (iii) where there are enough instances of convergence to allow sufficient statistical power. Here, we use the morphological diversity of stick and leaf insects (order Phasmatodea, ∼3,400 species) to provide such test. Most species exhibit stunning forms of camouflage through background matching (crypsis (28)) and the mimicry of objects irrelevant to predators (masquerade (29)) such as sticks, leaves, bark pieces, or moss (30–32). Selection to match such diverse objects produced a spectacular morphological diversity ranging from elongated tubular bodies with long slender legs to bodies so wide and flattened they look like leaves (Fig. 1). Recent phylogenetic studies of phasmids conflict with prior taxonomic classifications based on morphological characters, suggesting a high degree of morphological convergence across the Phasmatodea (33–37). For example, the “tree lobsters” – flightless, robust and strongly armored species, including the famous Lord Howe Island stick insect—had been grouped into the subfamily Eurycanthinae but were later shown to be highly polyphyletic, illustrating a dramatic case of morphological convergence (33) (Fig. 1 O,P). Moreover, several authors have suggested that apterous, stockier, spinier, and darker body forms tend to be found close to the ground, while more elongated and winged forms tend to rest higher up in the vegetation, implicating a role of ecological niche in driving these convergent patterns (31, 38). We quantitatively assessed the presence and extent of convergent evolution in body morphology in stick insects using a time-calibrated multilocus phylogeny of the order and an associated morphospace of female body morphology. Our analyses identified 20 distinct body types (ecomorphs), many described here for the first time, and revealed dozens of instances of morphological convergence, as behavioral transitions towards similar habitat uses were associated with the invasion of restricted and distinct portions of the morphospace. Using the two most common habitat transitions (15 and 14 transitions, respectively), we then examined how divergence time, ancestral habitat, and environmental distance affected the extent of morphological convergence.

**Figure 1:**
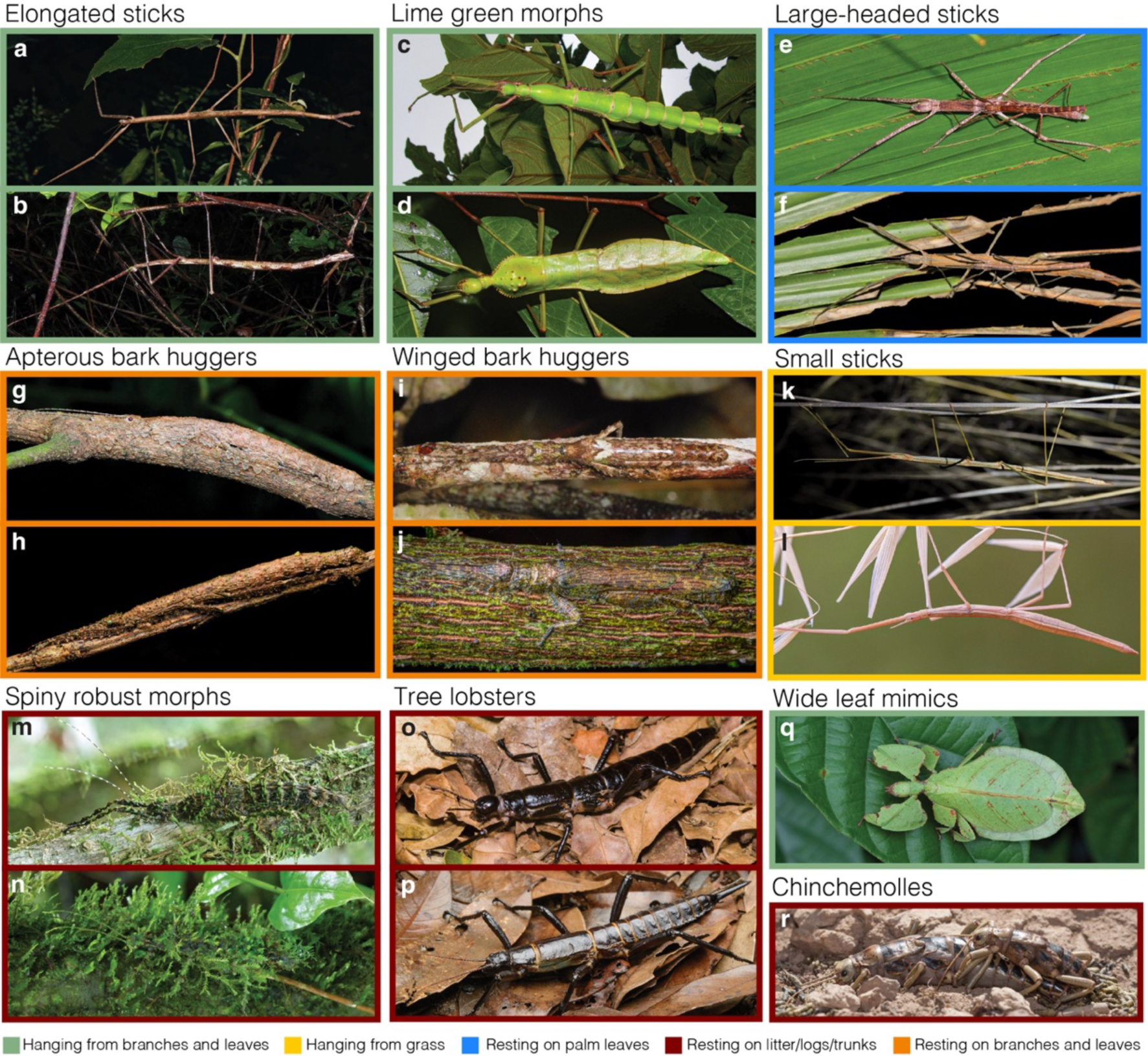
Photographs of adult females *in situ*. The color surrounding each picture correspond to a habitat use category. Pictures included under the same ecomorph name represent cases of convergent evolution (i.e., unrelated lineages). **a,** *Ctenomorpha marginipennis* (Australia, Lanceocercata) (CC-BY-NC 4.0 Julie Graham, inaturalist.org/observations/73831515); **b,** *Phobaeticus kirbyi* (Malaysia, Pharnaciini) (CC-BY-SA 2.0 Bernard Dupont, flickr.com/photos/berniedup/8419620988). **c,** *Monandroptera acanthomera* (Réunion, Lanceocercata) (© Nicolas Cliquennois, used by permission); **d,** *Cranidium gibbosum* (French Guiana, Diapheromerinae) (CC-BY-NC 4.0 Sébastien Sant, inaturalist.org/observations/75953936). **e,** *Apterograeffea reunionensis* (Réunion, Lanceocercata) (© Nicolas Cliquennois, used by permission); **f,** *Graeffea crouanii* (French Polynesia, Lanceocercata) (CC-BY-NC 4.0 Tahiticrabs, inaturalist.org/observations/165663078). **g,** *Leosthenes aquatilis* (New Caledonia, Lanceocercata) (CC-BY-NC 4.0 Damien Brouste, inaturalist.org/observations/24180348); **h,** *Pseudoleosthenes irregularis* (Madagascar, African/Malagasy clade) (© Paul Bertner, used by permission). **i,** *Epicharmus marchali* (Mauritius, Lanceocercata) (© Sylvain Hugel and Nicolas Cliquennois, used by permission); **j,** *Prisopus berosus* (Belize, Pseudophasmatinae) (CC-BY-NC 4.0 Thomas Shahan, inaturalist.org/observations/50919578). **k,** *Denhama sp.* (Australia, Lonchodinae) (CC-BY-NC 4.0 Enot Poluskuns, inaturalist.org/observations/166373254); **l,** *Clonopsis gallica* (Spain, European clade) (CC-BY 2.0 Ramón Portellano, flickr.com/photos/118276383@N05/14324421727/). **m,** *Parectatosoma sp.* (Madagascar, African/Malagasy clade) (© Paul Bertner, used by permission); **n,** *Taraxippus samarae* (Panama, Cladomorphinae) (© Paul Bertner, used by permission, inaturalist.org/observations/19995010). **o**, *Dryococelus australis* (Australia, Lanceocercata) (© Angus McNab, used by permission); **p**, *Eurycantha immunis* (Papua, Indonesia, Lonchodinae) (© Chien C. Lee, used by permission). **q**, *Pulchriphyllium bioculatum* (Singapour, Phylliidae) (CC-BY-NC 4.0, Catalina Tong, inaturalist.org/ observations/154447000); **r**, *Agathemera crassa* (Chile, Pseudophasmatinae) (CC-BY-NC-SA 4.0 Ariel Cabrera Foix, inaturalist.org/observations/29411794).

## Results

### Repeated evolution of ecomorphs in the Phasmatodea

We used genetic data (from 3 nuclear and 4 mitochondrial genes) from 314 phasmid taxa and fossil-calibrated Bayesian inferences to reconstruct the evolutionary history of Phasmatodea. The relationships between the major euphasmatodean clades that arose during an ancient radiation were constrained to match the basal topology inferred in previous phylogenomic studies (35, 39). The inferred Maximum Clade Credibility (MCC) tree was overall strongly supported and was largely congruent with previous studies (Fig. 2)(34, 37, 40), providing a robust framework for all subsequent comparative analyses. 16 major clades were recovered and appeared largely defined by geographic distribution and ecozones (Fig. 2). The split between Embioptera and Phasmatodea is estimated to have occurred 125 million years ago (mya) [95% Highest Posterior Density (HPD): 122 – 130mya] and between Timematidae and Euphasmatodea to 102mya [95% HPD: 99 – 108mya].

**Figure 2:**
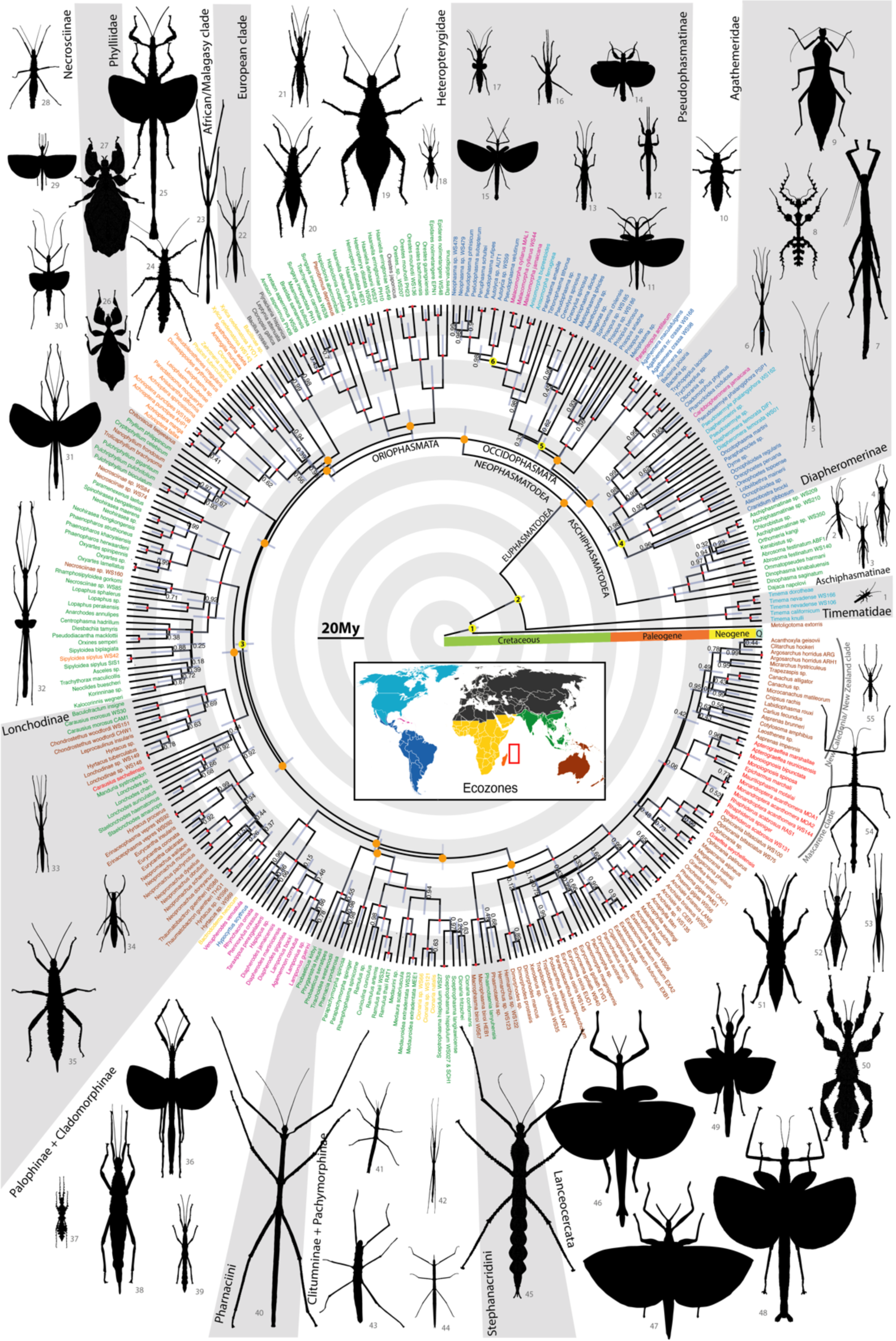
Time-calibrated maximum clade credibility tree and geographic distribution of stick and leaf insects. Fossil calibration points are denoted with numbered yellow circles (Table S1). Orange circles correspond to constrained nodes based on the topology of Tihelka et al. (2020). 95% confidence intervals around node ages are indicated by gray bars and Bayesian posterior probabilities are indicated at each node. Red nodes represent fully supported nodes with posterior probabilities equal to one. Tip labels are colored by ecozone following the colors of the central inset. The red rectangle on the world map indicates islands of the Mascarene plateau. Scaled adult female silhouettes were drawn by the first author and correspond to the species listed in Table S2.

We assembled a morphological dataset comprising more than 1300 adult female specimens from 212 species included in the phylogeny and including 23 quantitative measurements and qualitative data on color and body texture (Fig. S1). From this dataset, we reconstructed a multidimensional morphospace using a mixed Principal Component Analysis (PCAmix). The PCA revealed large variation between phasmid species in relative body width (PC1, 44% of the total variation), relative body height (i.e., how flattened the body is; PC2, 11%), relative wing size (PC3, 11%), body texture (i.e., how smooth or rough the body cuticle is; PC4, 8%) and relative head size (PC5, 5%) (Fig. 3A-B, S2). The first five PCs together accounted for 77% of the total variation. When considering the two first axes (accounting for 55% of the total variation), morphological variation in phasmids appeared bimodal between the wide, leaf-mimicking Phylliidae clade and the rest of the phasmids, which display a more stick-like, tubular morphology (Fig. 3A). Phylliids, often referred to as true leaf insects, are characterized by an exceptionally widened and flat abdomen giving them the appearance of wide angiosperm leaves (Fig. 1Q) (41). All the other phasmid clades appeared centered around a single core on the morphospace, varying mostly in relative body width ranging from extremely elongated to more robust body silhouettes (Fig. 3A). Species with extreme morphologies were scattered at the periphery of this central core, often only projecting out along a single axis. For instance, the large-headed palm stick insects (subfamily Megacraniinae) mostly stand out along the PC5 axis that separates species based on relative head size (Fig. 1 E,F). Most of the morphological diversity is found in the Euphasmatodea, consistent with their much greater species diversity (n>3400 species), compared to Timematodea (n=21 species), which is morphologically homogeneous (Fig. 3). The reconstructed explosive morphological diversification of Euphasmatodea can be visualized in Movie S1.

**Figure 3:**
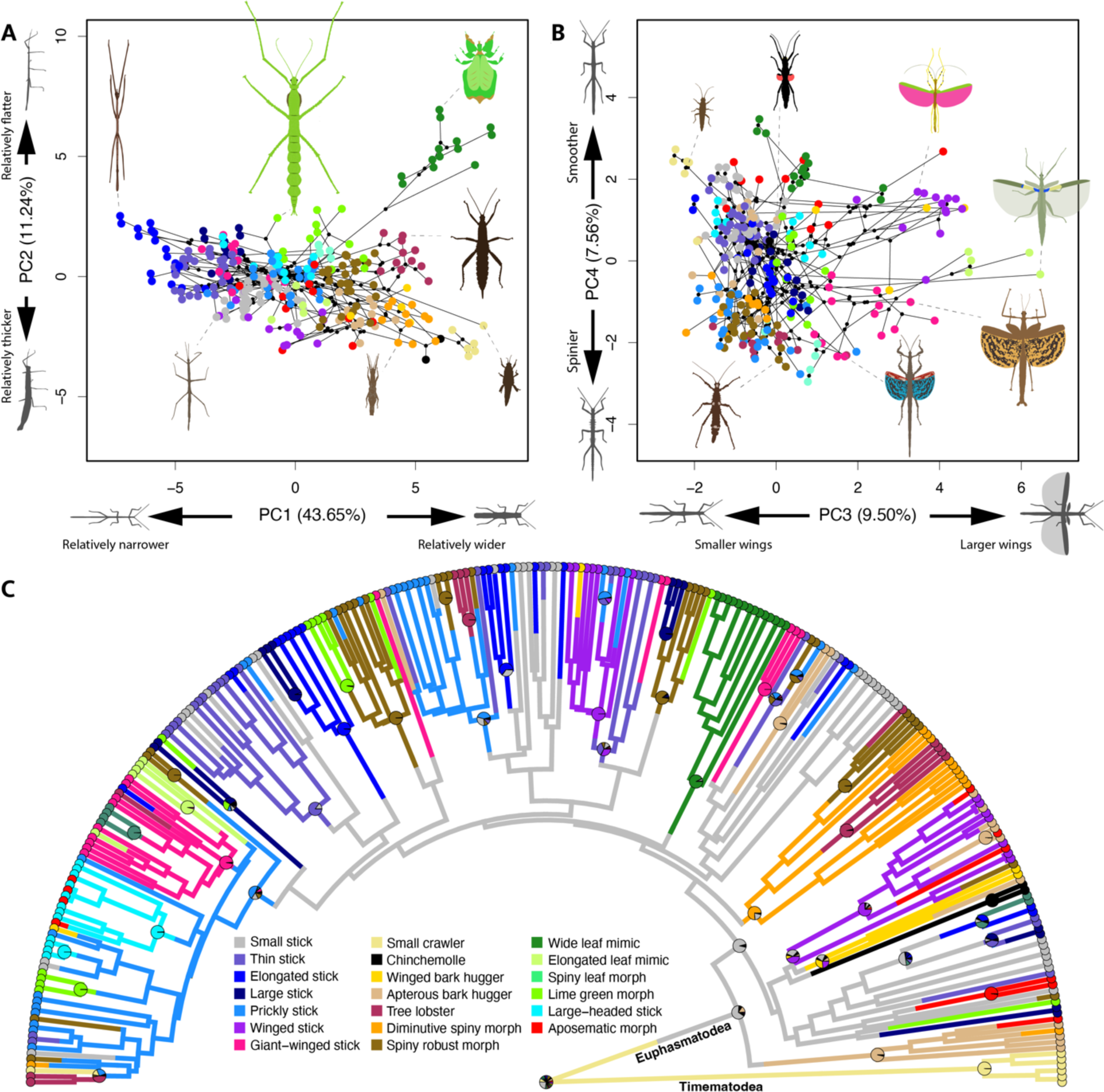
Ecomorphological convergence in stick and leaf insects. **A-B:** Phylomorphospace (first four dimensions) with species colored by assigned ecomorph (see fig. 4). Black lines between points represent the underlying phylogenetic relationships between species. **C**: Ancestral state reconstruction of ecomorphs using stochastic character mapping. The pie charts at nodes represent the posterior probabilities that each internal node is in each state. The color legend applies to all panels.

We then used a hierarchical clustering approach to define and assign species to clusters occupying relatively distinct regions of the multidimensional morphospace (Fig. 3A-B, 4, S3-5). The optimum number of clusters (k=20) was determined using the biological homogeneity index [BHI, (42)] to maximize the homogeneity of habitat use within each cluster (Fig. S6). BHI measures the average proportion of taxa pairs with similar habitat uses and which are clustered together morphologically. For 20 clusters, BHI was 0.82, which highlights the strong association between habitat use and the defined morphological clusters, thereafter referred to as ecomorphs. As expected, among the 20 ecomorphs, we recovered the *wide leaf mimic* ecomorph (Fig. 1Q) and the previously recognized *tree lobster* ecomorph (Fig. 1 O,P), which includes the thorny devil stick insects (*Eurycantha* spp.) and the Lord Howe Island stick insects (*Dryococelus australis*)(33). Using random forest machine learning models, we identified the main morphospace axes that were most helpful for these predictive models to infer ecomorph from the morphological data and therefore the axes best distinguishing each ecomorph (Table S4). This analysis revealed that ecomorphs are often distinguished by only a few dimensions of the morphospace. For instance, *spiny robust morphs* were best distinguished by PC1 (i.e., relative body width) and PC4 (i.e., body texture) due to their stocky and spiny bodies, often mimicking bark pieces or moss (Fig. 1 M,N; Table S4, Fig. S3).

**Figure 4:**
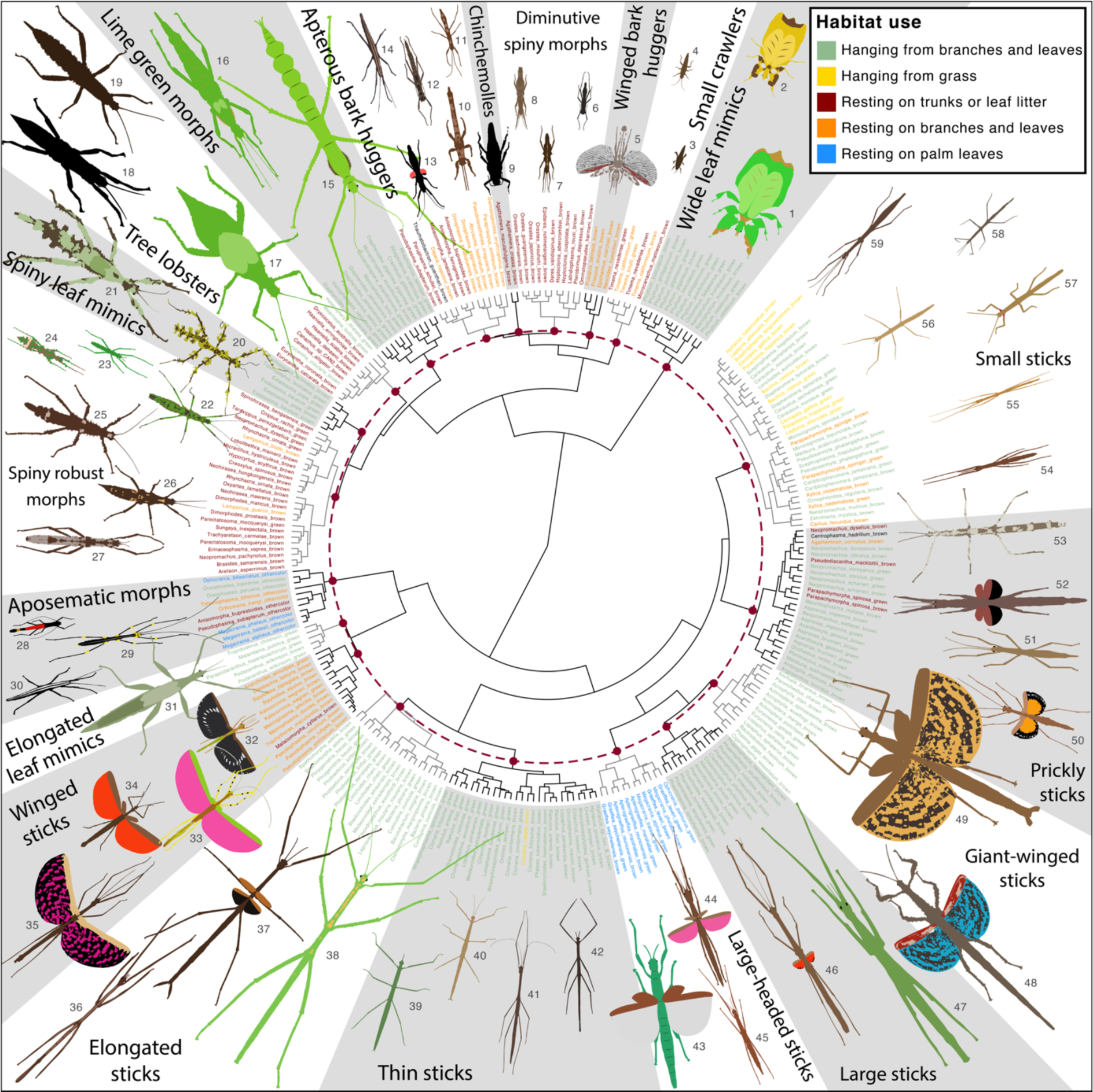
Phenogram of overall morphological similarity across female phasmids. Hierarchical cluster dendrogram based on 24 continuous and 3 discrete morphological variables using the Ward’s method. Tip labels are colored according to extant habitat use. The dashed maroon circle corresponds to the height threshold used to delineate ecomorphs. Intersection between the circle and dendrogram branches are shown as maroon dots. Scaled adult female illustrations were drawn by the first author and correspond to the species listed in table S3.

A discrete ancestral state reconstruction based on stochastic character mapping suggested that the *small stick* ecomorph (Fig. 1 K,L), characterized by its central position on the morphospace, was the ancestral ecomorph of Euphasmatodea (Fig. 3C). It further revealed that the *chinchemolles* (clade Agathemeridae) and the *wide leaf mimic* (clade Phylliidae) ecomorphs (Fig. 1 Q,R) each had unique origins and were therefore evolutionary one-offs. *Chinchemolles*, as originally named by native South Americans, are fat, robust and smooth insects adapted to the arid slopes of the Andes range and known for their foul defensive smell (from the Quechua chinche: “fetid”, and moyo: “udder”) (43, 44) (Fig. 1R). All other ecomorphs appeared to have originated at least twice (e.g., *small crawler* ecomorph) and up to at least 10 times (e.g. *thin stick* ecomorph), indicating widespread morphological convergence in the order (Fig. 3C).

### Phasmid morphology and habitat use are closely associated

A stochastic character mapping of habitat use reconstructed the ancestor of all phasmatodeans as most likely having rested on the leaf litter, trunks, or logs during the day (Fig. 5A). However, the ancestors of most euphasmatodean clades were inferred as hanging from branches and leaves (Fig. 5A). Overall, this reconstruction indicated between 15 and 19 secondary transitions to resting on the leaf litter, logs and trunks, 16 transitions to resting on branches and leaves, five to hanging from grass, and two to resting on palm leaves (Fig. 5A). We calculated the size of the multidimensional hypervolumes occupied by each habitat category on the morphospace using range boxes and kernel density estimates (45, 46). Species hanging from branches and leaves occupied the largest volume on the morphospace, species hanging from grass or resting on palm leaves the smallest (Fig. 5B, S7-8). This reflects the considerable variation in body morphology of species hanging from branches and leaves going from extremely elongated and cylindrical stick-like species (e.g., Fig. 1 A,B) to wide and flat leaf-like species (e.g., Fig. 1 Q) and wingless to fully winged species (e.g., Fig. 1 A-D, Q). Hypervolume overlap, as measured by different methods, was overall relatively low between habitat categories (Jaccard similarity ranged from 7.5e^-6^ to 0.17, Sorensen similarity from 1.5e^-5^ to 0.33) (Fig. S9-10). Random forest models (i.e., machine learning classification algorithms) reached 85.2% accuracy when classifying the habitat use of taxa based solely on morphospace coordinates (Fig. 5C-D). The accuracy of predictions was limited when only based on the first morphospace axis, despite PC1 accounting for almost half of the phenotypic variance (44%, Fig. S2), but plateaued after including the first 5 axes only (Fig. 5C).

**Figure 5:**
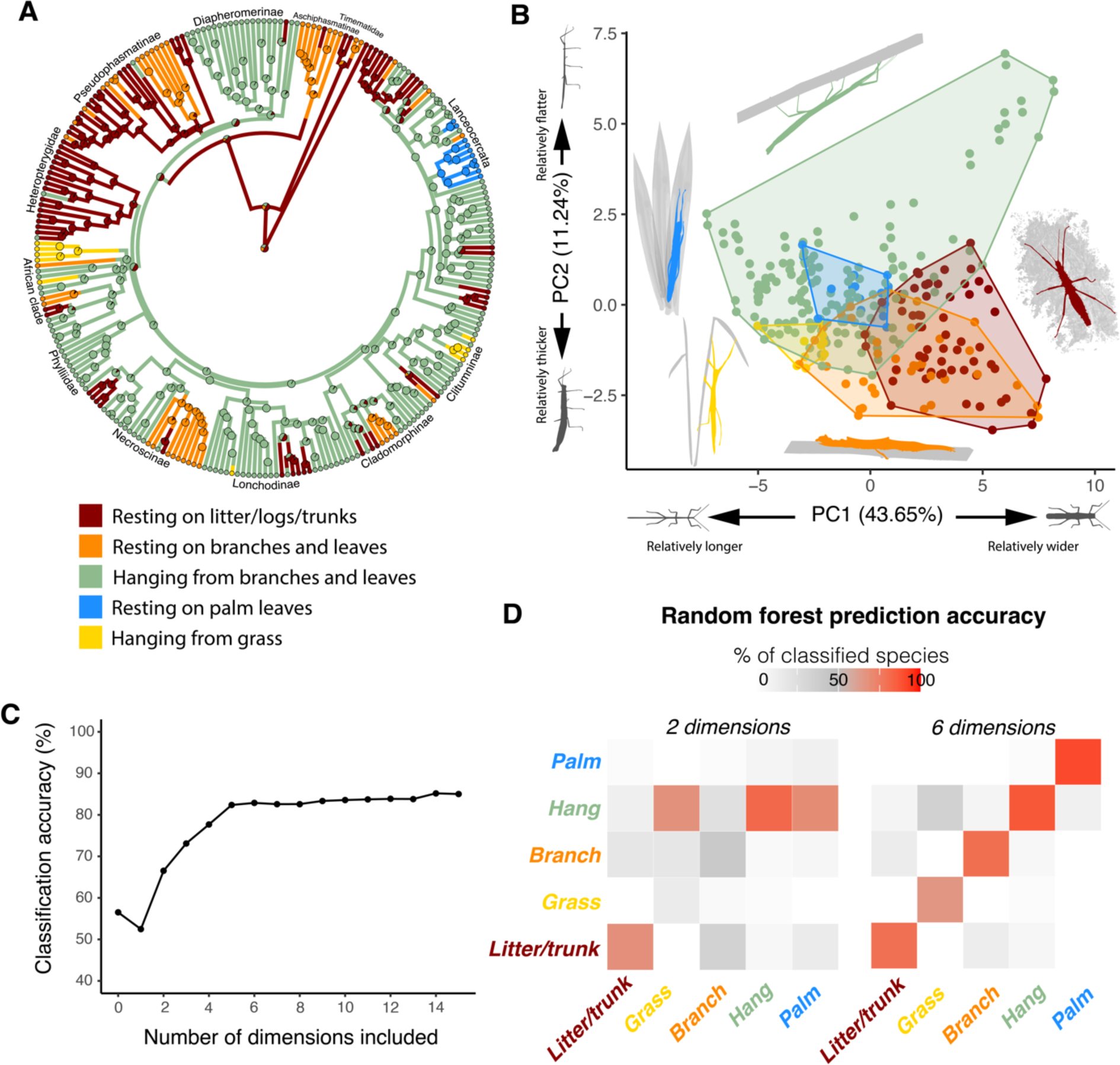
Habitat transitions and morphospace occupation and overlap between different habitats. **A:** Ancestral state reconstruction of habitat use using stochastic character mapping. **B**: Convex hulls of species sharing the same habitat use on the morphospace. **C**: Mean accuracy of the random forest model at predicting habitat use based on the number of morphospace axes provided. **D:** Heatmaps showing the prediction accuracy of random forest models for each habitat use based on two or six morphospace axes. Predicted habitat states are displayed on the x axis and observed habitat states on the y axis.

### Habitat transitions are associated with parallel shifts towards the same morphospatial regions

We used a series of complementary process- and pattern-based approaches to quantitatively assess the strength of morphological convergence between lineages independently transitioning towards the same habitat category (thereafter called “convergent lineages”). First, we compared the relative fit of a set of multivariate models of trait evolution [mvMORPH, (47)] and found support for the multi-regime Brownian motion model (BMMm), with distinct regimes corresponding to the five different habitats (Table S5). Thus, habitat use affects morphological evolution but does not correspond to a unique optimum as BMM models do not model selection toward optima, unlike Ornstein–Uhlenbeck (OU) models which provided worse fits of the data (Table S5). In contrast, when considering different regimes for the different ecomorphs instead of habitat categories, the best supported model was by far the multi-regime OU model (Table S6). Therefore, ecomorphs could correspond to adaptive optima as they occupy much smaller regions of the morphospace.

We then assessed the phenotypic similarity between convergent taxa and distinctiveness from other taxa [Wheatsheaf index (*w*), (48, 49)] and the increase in similarity between the convergent taxa through time [C1 to C4 metrics (C-metrics), (9)]. *w* identified significantly stronger convergence for lineages that independently transitioned to resting on the ground/trunks, to resting on branches, and to hanging from grass than would be expected from a random distribution of trait values simulated under a Brownian Motion (BM) model (P < 0.02, Table 2). Likewise, most of the C1 to C4 statistics were significantly higher than expected under random evolution for all habitats except lineages secondarily transitioning back to hanging from branches (Table S7).

The C-metrics rely on the difference between the present distance on the morphospace between two convergent lineages (D_tip_) and the maximum distance between any two points (not necessarily synchronous) along the evolutionary trajectories of the two lineages (D_max_, Fig. 6C). Consequently, these metrics can be equally high for lineages that had very dissimilar ancestors at some point in time but that subsequently became more similar, and for lineages shifting in parallel towards a similar region of the morphospace (50). To distinguish between these two scenarios, we computed the recently developed C_t_ measures, which compare the extant phenotypic distance between the convergent lineages to the maximum reconstructed ancestral distance at a given time point during their evolution (i.e., between synchronous points along the evolutionary trajectories)(50). Unlike C-metrics, C_t_-metrics are only expected to be high when lineages moved away from one another at some point in their evolutionary history and then subsequently got closer (i.e., converged). C_t_ measures were only significantly higher than expected by chance for transitions to resting on the leaf litter or trunks (Table S7). But even in this case, C_t_ values were relatively close to zero or even negative, indicating that convergent taxa are not necessarily morphologically closer to one another than their ancestors. This suggests that lineages independently evolving similar habitat uses shifted in parallel towards the same broad region of the morphospace, and sometimes even diverged in that novel region (i.e., “imperfect” convergence (51)) (Fig. 6A-B). Parallelism was further confirmed by calculating the pairwise angles between the evolutionary trajectories of convergent lineages following the independent invasion of a given habitat (θ, Fig. 6C). θ was lower than expected by chance for all habitat transitions, except for secondary transitions to hanging from branches (Table S7).

**Figure 6:**
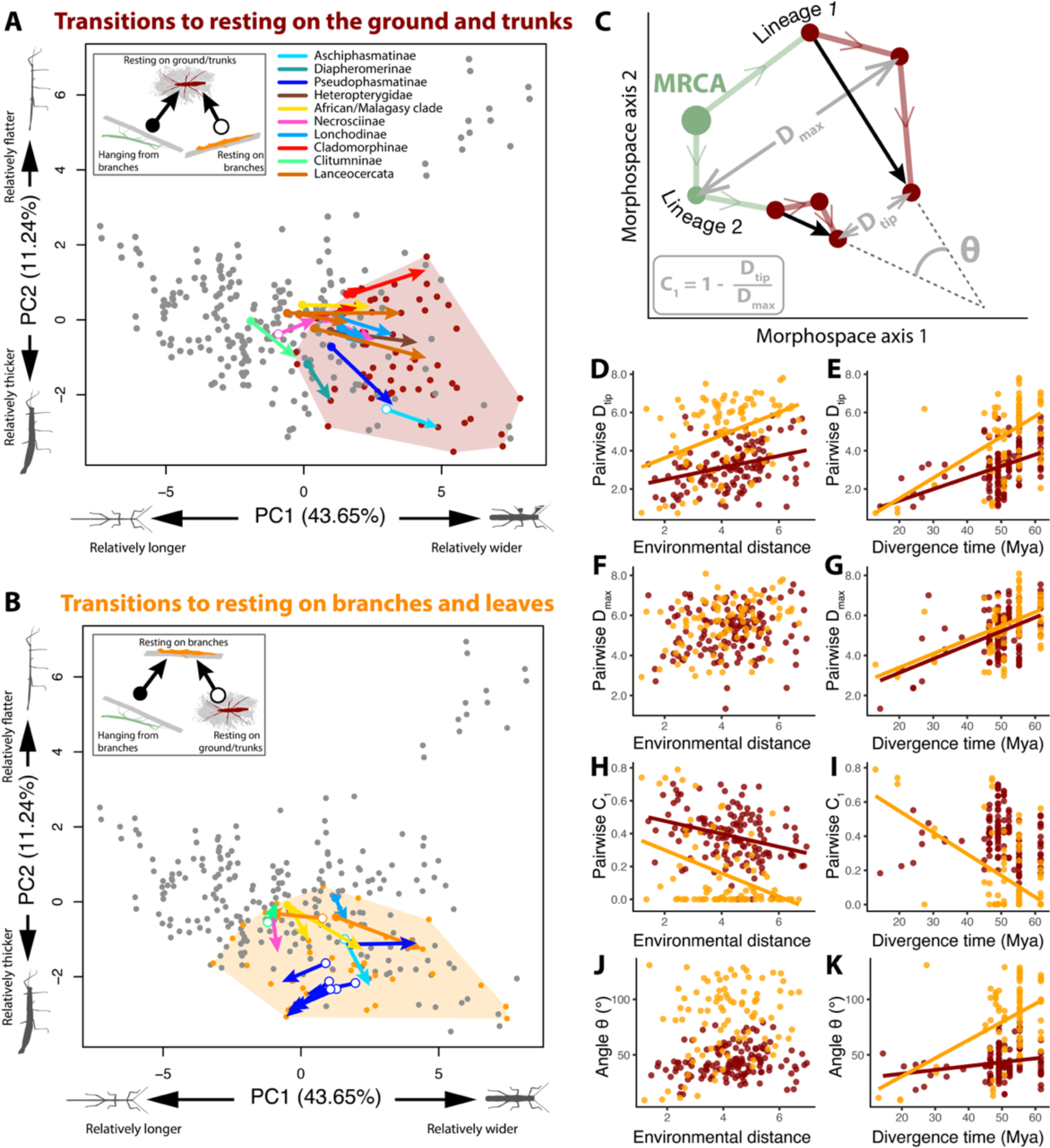
Evolutionary trajectories and effects of environmental distance and divergence time on morphological convergence. **A-B:** Trajectories on the morphospace of lineages that independently transitioned to resting on the ground and trunks (maroon, **A**) or to resting on branches and leaves (orange, **B**). Corresponding convex hulls are shown. Arrows start at the inferred position of the ancestor that first transitioned to the new habitat. Arrows end at the centroid position of descendant species. Arrow colors correspond to genetic clades (see fig.1). Start symbols indicate the ancestral habitat from which each lineage transitioned according to the insets. **C.** Example of the calculation of measures of convergence. The independent trajectories over time of two lineages splitting from their Most Recent Common Ancestor (MRCA) are shown on a morphospace. These lineages start as hanging from branches (pale green) and independently transition to resting on the ground and trunks (maroon). Circles represent ancestral nodes or tips. D_tip_ shows the current morphological distance between the two tips of interest. D_max_ shows the maximum distance between the lineages at any point in time between the tips and the MRCA. C_1_ calculates the proportion of the maximum distance between two lineages that has been erased by convergent evolution (0 ≤ C_1_ ≤ 1). θ represents the angle between the two vectors starting from the first nodes in the new habitat state and ending at the tips. It compares the overall direction of change between lineages after independently invading the same habitat. **D-K**: Pairwise D_tip_ (**D-E**), pairwise D_max_ (**F-G**), pairwise C_1_ (**H-I**) and pairwise θ (**J-K**) as a function of pairwise environmental distance and pairwise divergence time for each independent transitions to resting on the ground and trunks (maroon) and resting on branches and leaves (orange). Linear regressions are only shown if the effect of environmental distance and divergence time on the response variable are significant (see Table S8).

### Environmental similarity and phylogenetic relatedness promote stronger morphological convergence

Whether two lineages converge toward closer morphospatial regions following a similar habitat transition may be affected by several factors including whether they started from the same ancestral habitat, the extent of environmental similarities between their new habitats, and their phylogenetic relatedness. We took advantage of the repeated independent transitions toward resting on the leaf litter and trunks (n=15, Fig. 6A) and towards resting on branches and leaves (n=14, Fig. 6B) to test the effect of these three factors on the strength of convergence. Other transitions were too rare to allow sufficient statistical power (n ≤ 4, Fig. 5A). We calculated pairwise environmental distances between lineages as the distance on a multidimensional environmental space built from various macroecological variables relating to habitat height, climatic conditions, plant productivity and predator diversity (Fig. S11). For each pair of lineages, we also scored whether they transitioned from the same habitat state or not (binary) and their divergence time as the age of their most recent common ancestor. Convergence between pairs of lineages was quantified as pairwise D_tip_, D_max_, C_1_ and θ (Fig. 6C).

For both habitat types, multiple matrix regressions revealed that environmentally closer lineages (i.e., lineages colonizing more similar selective environments) and more closely related lineages transitioned toward closer positions on the morphospace (i.e., lower D_tip_; Fig. 6D-E, Table S8). D_max_ was only significantly affected by divergence time between the lineages: lineages that diverged a long time ago are more likely to exhibit a large D_max_ relative to more closely related ones (Fig. 6F-G, Table S8). Consequently, C_1_ decreased with environmental distance indicating weaker convergence between lineages experiencing more dissimilar environmental conditions (Fig. 6H, Table S8). C_1_ was negatively affected by divergence time but only significantly so in lineages that transitioned to resting on branches and leaves (Fig. 6I, Table S8). θ was only affected by divergence time: more related species tend to follow more parallel evolutionary trajectories (Fig. 6J-K, Table S8). Lineages that transitioned toward the same habitat from different ancestral habitats and thus potentially starting from further apart on the morphospace exhibited less parallel trajectories. However, the significance of this effect was low (p≥0.09, Table S8).

## Discussion

When adapting to shared environmental challenges, lineages often vary in the extent to which they evolve similar traits, indicating that evolutionary outcomes are more predictable in some instances than in others. Explaining this variation will be critical as scientists increasingly base medical (vaccine design, pandemic preparedness, antibiotic resistance, cancer therapies), agricultural (application of herbicides and pesticides, anticipating crop responses to climate change), and conservation (wildlife responses to anthropogenic disturbance and climate change) practices on predicted evolutionary responses to selection (52, 53). Here we used the dozens of repeated instances of convergent phenotypic evolution in stick and leaf insects to quantify the relative contributions of divergence time (phylogenetic relatedness), similarity of most recent ancestor habitats, and the similarity of invaded habitats, on the repeatability – and therefore the predictability – of evolutionary outcomes. As in earlier studies of repeated evolution, we show that closely related lineages (i.e., likely sharing more genetic variation) followed more parallel evolutionary trajectories and ended up relatively closer on the morphospace, consistent with the idea that in the absence of opportunity for contingency, phenotypic responses to selection will be highly predictable (27, 54). However, our study encompassed a wide range of divergence times (10 to 60 million years), and we also show that the strength of morphological convergence decreases steadily with time since divergence (Fig. 6D-K, Table S8). Ironically, this suggests that the stochastic contributions of contingency are also predictable, in the sense that they accrue at a constant rate over time.

Classic examples of morphological convergence are often found among closely related taxa (e.g., *Anolis* lizards (14, 15), stickleback fish (16), cichlid fish (55)), suggesting that the repeatability of phenotypic evolution increases with relatedness (10). Here we provide an original and direct test of this idea in a system spanning vast divergence times (10 to 60 million years)(35, 39). Closely related lineages appear predisposed to adapt in more similar ways when confronted with similar challenges, consistent with Gould’s idea that evolution is less inclined to repeat itself at large macroevolutionary time scales (17). The higher level of repeatability of morphological evolution observed among closely related lineages manifests itself in two ways: more closely related, convergent lineages look more similar, and also followed more parallel paths on the morphospace to arrive there. Closely related lineages share more standing genetic variation and segregating variants are expressed against more similar genetic backgrounds (21, 56–58). And they are more likely to reuse the same genes when they adapt to similar environmental challenges (22–25).

The other factor likely to influence the strength of convergence is the environment: the more similar the selective conditions experienced by two lineages, the closer the resulting convergent phenotypes. Studies of phenotypic convergence often categorize ecological niches to identify associations between patterns of morphological evolution and the repeated adaptation to these discrete niches (e.g., diet types (51), lakes/streams (6, 12)). This categorization hides potential heterogeneities in environmental conditions among instances of the same category. Conditions that appear similar to a human observer may actually be disparate to the organisms, and this can confound studies attempting to explain variation in the strength of convergence. For example, stickleback fish independently colonizing stream habitats varied in the extent of their phenotypic convergence in part because habitats categorized as “stream” actually differed in water clarity, temperature, parasite abundance, and food availability (27). Once these additional variables had been included, habitat similarity predicted the resulting strength of convergence more accurately (27).

Here we quantified habitat similarity using various macro-ecological variables, and our results suggested that some of the niches invaded by phasmids (e.g., grass) were environmentally restricted and thus very similar across lineages, while others (e.g., resting on branches and leaves) encompassed much wider and potentially less similar habitat conditions (i.e., they likely included “cryptic” habitat dissimilarities) (Fig. S12). We show that similarity of invaded habitats also predicted strength of convergence, such that lineages switching to more similar environments ended up in closer regions of the morphospace (Fig. 6D,H). Thus, our study highlights that this effect also applies across large macroevolutionary scales, despite higher levels of contingency.

Variation in the specific environmental conditions experienced by each lineage within the same habitat category may explain, at least partly, why these categories correspond to broad regions of the morphospace. Consistently, the best model for the evolution of phasmid morphology is a Brownian motion model with different habitat categories corresponding to different mean values and evolutionary rates (Table S5). The lack of support for Ornstein– Uhlenbeck models, which explicitly model selection toward adaptive optima, highlights that a given habitat category does not correspond to a fixed adaptive optimum. Instead, the position of adaptive optima may be dependent on the specific environmental conditions uniquely experienced by each lineage. Alternatively, or additionally, habitat categories may reflect different rugged adaptive landscapes with multiple peaks corresponding to the various ecomorphs identified. The exact position of these peaks may be altered by lineage-specific environmental conditions. Phasmids, which mainly rely on camouflage for predator defense, depend on substrate appearance for the efficiency of crypsis and on the surrounding items’ diversity for masquerade. Thus, it would make sense that a habitat category like “resting on branches” would have an adaptive landscape with multiple fitness peaks, each corresponding to a different type of branch.

Finally, we accounted for similarity of the most recent *ancestral* habitats of convergent pairs of lineages, to test whether transitioning from the same or different habitats affected the extent of the resulting convergence in this group of insects. Lineages that converged from the same ancestral habitat tended to follow more parallel trajectories, but this effect was only marginal (Table S8). It is possible that there were not enough transitions from the same versus different ancestral habitats to detect this effect.

The Euphasmatodea show a deep radiation at the base of the group (∼65–55Mya) following the K-T boundary (35), corresponding with the origin of most major clades and with dispersal across vast regions of the globe (Fig. 2). Although a few of these clades seem to have undergone speciation without niche differentiation, and species within these clades are morphologically homogeneous (e.g., Phylliidae (wide leaf mimics and canopy-dwellers) and the Heteropterygidae (spiny and robust ground-dwellers), which are distributed on many islands of Indomalaya and Australasia (Fig. 3C, 5A) (40, 41)), the majority of euphasmatodean clades subsequently radiated into multiple different ecomorphs colonizing diverse habitats (e.g., Lanceocercata (Australasia and Mascarene islands), Cladomorphinae (Caribbean islands), Lonchodinae (Indomalaya/ Australasia), Necrosciinae (Indomalaya), African/Malagasy clade (Afrotropics), Pseudophasmatinae (Neartic and Neotropics) and Diapheromerinae (Neartic and Neotropics); Fig. 3C, 5A) (33, 40, 41, 59). We characterized 20 different phasmid ecomorphs and reconstruct dozens of evolutionary transitions between ecological niches, resulting in repeated instances of convergence towards these phasmid body forms. Overall, our results suggest the extremely diverse morphologies of stick and leaf insects result from replicated radiations in different geographic regions, each associated with widespread parallel shifts on the morphospace as independent lineages adapted to similar habitats.

## Conclusion

Phasmids exemplify the extraordinary power of natural selection to shape organismal phenotypes. The animals themselves are charismatic champions of crypsis and masquerade, and our comprehensive quantification of their trajectories of evolution, using process-based (i.e., evolutionary modelling) and pattern-based methods, reveals dozens of instances of convergence. Our findings show that the details of the environmental conditions experienced by the organisms – the closeness of the invaded niches – predicts the extent of convergence even when lineages have been evolving independently for tens of millions of years, and therefore have had ample opportunity for contingency. Furthermore, we show that the effects of contingency also are relatively predictable, gradually eroding the strength of convergence at a steady rate across spans of up to 60 million years. We suggest that precise quantification of selective environments, as well as divergence times, will be critical as studies increasingly attempt to predict the outcomes of evolution.

## Materials and Methods

Extended materials and methods are reported in the SI Appendix, Supplementary Materials and Methods, and include details on definitions and choices of convergence metrics.

### Taxonomic sampling and phylogenetic reconstruction

Well-supported phylogenies for 38 phasmid lineages representing all major clades of Phasmatodea were recently reconstructed using next-generation sequencing (transcriptomes), yielding topologies that resolved most of the deep nodes within this group with high confidence (35, 39). Here we reconstructed a phylogeny with 314 species representing all major phasmid lineages (9% of the known phasmid species diversity and 33% of currently recognized generic diversity), and one species of Embioptera (the sister clade of Phasmatodea) as outgroup, constraining the basal topology to match the transcriptome-based trees (39). Regions of 3 nuclear (18S rRNA (18S), 28S rRNA (28S) and histone subunit 3 (H3)) and 4 mitochondrial genes (12S rRNA (12S), 16S rRNA (16S), cytochrome-c oxidase subunit I (COI) and cytochrome-c oxidase subunit II (COII)) were extracted from Genbank, aligned and concatenated (6,778bp total) to reconstruct a Maximum Clade Credibility (MCC) tree for phasmids using Bayesian inferences in BEAST 2 (v. 2.6.3)(dataset S1)(60). Divergence time was estimated using 6 unambiguous crown-group phasmid fossils as minimum calibration points (Table S1).

### Morphological data

We examined 1359 adult female specimens from 212 species included in the phylogeny. High-quality photographs, captured in dorsal and/or lateral views, were obtained from our own collection at the University of Göttingen (Germany), other museum collections, the published literature and other online sources (dataset S1). Depending on material availability, we measured pictures of between 1 and 18 different individuals per species (mean = 5.5 individuals per species). We collected 23 continuous measurements (Fig. S1) that together contained biologically relevant information about overall body size and shape, width and length of different body segments (notably the head), leg length, hindwing size and the length of the subgenital plate (whose function is often related to oviposition). We also qualitatively scored body texture and overall body coloration to include these critical aspects of camouflage when reconstructing the morphospace.

### The phasmid morphospace

We built a multidimensional morphospace using a Principal Component Analysis (PCA) mixing continuous and categorical data (PCAmix)(61). Continuous measurements were log_10_-transformed. Body volume and area were included as body size measurements. Then, we included the remaining traits after controlling for the effect of size by substituting original trait values with the residuals calculated from a phylogenetically-corrected linear regression against body volume. Because wing length and wing area included zeros for wingless species, we divided the non-transformed measurements by body length or body length squared respectively, to obtain and include measures of relative wing length and area. In total, we included 23 continuous (priorly mean-centered on zero and scaled to unit variance) and three categorical variables (Fig. S1).

### Habitat data

We broadly classified the habitat of stick insects based on the typical resting posture and substrate preferences exhibited by adult females when hiding during the day (i.e., when they are exposed to visually hunting predators). We surveyed the literature, field guides and iNaturalist (https://www.inaturalist.org/, accessed July 2021) for observations of where each species is typically found (dataset S1). We defined five habitat use categories: resting on the ground or trunks (including the base of trunks, mossy logs, under bark, in the leaf litter), resting on branches and leaves, hanging from branches and leaves, hanging from grass, and resting on palm leaves. It should be acknowledged that this classification is broad and consequently does not fully encompass the entire spectrum of substrates and host plants upon which phasmids may be found (30–32).

### Environmental data

We gathered information about the geographic range of each species based on sampling location of type specimens and observations on iNaturalist (available from https://www.inaturalist.org, accessed July 2021). For each species, we then selected the median location with the most central latitude. From the GPS coordinates of the most central location for each species, we extracted data on 17 environmental variables that together contained information about climatic conditions (temperature, precipitation, seasonality), vegetation density and food availability (primary production), predator diversity and habitat vegetation layer (see Supplementary information, dataset S1). Variation in these variables was summarized by running a principal component analysis (Fig. S11).

### Definition of ecomorphs

We used our multidimensional morphospace data to cluster species into distinct ecomorphs by running a hierarchical clustering algorithm (using the Ward’s method) to define ecomorphs based on overall proximity on the morphospace (defined by the first 15 PC axes). We defined the optimal number of clusters thanks to the computation of a Biological Homogeneity Index (BHI) which measured how homogeneous clusters are, based on habitat use (R package “clValid”) (42, 62). Clusters were defined by a fixed height threshold on the clustering dendrogram. The optimal number of clusters was then chosen as the minimal number of clusters to maximize BHI (Fig. S6). We then identified the morphospace axes that best distinguished each ecomorph by training random forest models (R package “randomForest”) (63) to classify a taxon in either an ecomorph of interest or in a different one, given the first ten axes of the morphospace (Table S4).

### Overlap between habitat categories on the morphospace

To quantify morphospace occupation by species exhibiting different habitat uses (Fig. 5B), we estimated 6-dimensional hypervolumes (PC1-PC6, accounting for 81% of the total variation) using dynamic range boxes (R package “dynRB”)(45) and high-dimensional kernel density estimations (R package “hypervolume”)(46). Pairwise hypervolume overlap was quantified as the portion of the hypervolume of habitat A covered by the hypervolume of habitat B and vice versa, as the Jaccard similarity index (ratio of the intersection to the union of the hypervolumes), or as the Sørensen–Dice similarity index (ratio of twice the size of the intersection to the sum of the individual hypervolumes). The distance between the hypervolumes was also quantified as the Euclidean distance between the hypervolume centroids and the minimum Euclidean distance between points of the two hypervolumes. Finally, we also quantified the overlap between the habitat categories on the morphospace using machine learning random forest models (63). These models were used to predict the habitat category of a species given its position on the morphospace. The predictive error rate of the models was used to quantify overlap between habitat categories.

### Ancestral state reconstruction of ecomorphs and habitat use

Habitat use was mapped on the MCC tree to uncover the number of independent transitions toward each of the five categories. We ran ancestral state reconstructions using stochastic character mapping as implemented in the R package “phytools” (64). The transition matrix was calculated using maximum likelihood and using an all-rates-different model (model= “ARD”). Ecomorphs, as defined by our hierarchical clustering analysis, were similarly mapped to establish whether they had single or multiple origins. Given the large number of ecomorphs (n=20), only the “equal rate” transition model (assuming a single transition rate between ecomorphs) could be run.

### Process-based tests of convergence – evolutionary model fitting

To test for morphological convergence among lineages that independently transitioned to the same habitat, we fitted multivariate models of continuous trait evolution to PC1-PC5 (77% of the total variance) using the “mvMORPH” R package (47). We first fit the single-regime Brownian motion (BM1, modeling stochastic trait changes over time), Ornstein–Uhlenbeck (OU1, modeling stabilizing selection around an optimal trait value), and early burst models (EB, modeling stochastic changes with a decrease in evolutionary rate over time), which represent the null hypothesis. Then, for each habitat category, we fit 2-regime models where the given habitat category was considered its own evolutionary regime while the rest belonged to another unique regime. We also ran 5-regime models including each habitat as a separate regime, and 20-regime models including each ecomorph as a separate regime. The ancestral histories for each of the tested regime assignments were reconstructed on the MCC tree using 100 stochastic character maps (64). We fitted multi-regime OU models (OUM) allowing trait optima to vary among regimes, and BM models allowing on one hand the phylogenetic means to vary among regimes, and on the other hand holding the evolutionary rate constant (BM1m) or not (BMMm).

### Pattern-based tests of convergence

To quantify the strength of morphological convergence associated with repeated habitat transitions, we used the first five morphospace axes (PC1-PC5) to calculate the C1 to C4 pattern-based metrics (R package “convevol”)(9) as well as the Wheatsheaf index (*w*) (“windex”)(48, 49). C1-C4 are based on the ratio between the current distance between two lineages (D_tip_) on the morphospace to the maximum reconstructed distance between the two lineages at any point in the past (D_max_) (Fig. 6C). C1-C4 will be high when convergent lineages diverged substantially after splitting and subsequently re-evolved similarities, or when convergent lineages shifted in parallel towards the same direction on the morphospace (50). To distinguish between the two scenarios, we computed the recently developed Ct1-Ct4 metrics, which restrict D_max_ to synchronous nodes (50). Ct1-Ct4 are only expected to be high in the first scenario (divergence then convergence). Finally, we quantified parallelism in the evolutionary trajectories of convergent lineages by calculating the angle (θ) between these trajectories on the morphospace. We reconstructed the trajectories of convergent lineages from the position of the node immediately prior to the inferred habitat transition, to that of the tip of interest (Fig. 6C). For each above-described variable, the associated p-values test the hypothesis that convergence is significantly stronger (or that trajectories are more parallel) in the habitat category of interest than would be expected by chance.

### Analysis of the variation in the extent of morphological convergence

We tested the effects of three factors on the extent of morphological convergence: the phylogenetic relatedness between the convergent lineages, their environmental similarity, and whether they started from the same ancestral habitat. We only considered the repeated transitions toward resting on the leaf litter and trunks (n=15, Fig. 5A) and towards resting on branches and leaves (n=14, Fig. 5B) for these analyses as other transitions were too rare to allow sufficient statistical power (n ≤ 4, Fig. 4A). For each transition type, phylogenetic relatedness, environmental distance, ancestral habitat difference and morphological convergence were computed for all possible pairs of taxa corresponding to separate independent transitions toward the habitat category, and then assembled as distance matrices. Pairwise phylogenetic relatedness was estimated as the age of the most recent common ancestor of the two lineages. Pairwise environmental distance was calculated as the Euclidean distance on the environmental PC1-PC4 (accounting for 80.1% of the total environmental variation). Pairwise ancestral habitat difference was scored as either 0 if both lineages transitioned to the habitat of interest from the same ancestral habitat, or 1 otherwise. Finally, to quantify morphological convergence we computed pairwise D_tip_, pairwise D_max_, pairwise C_1_ and pairwise θ. We fitted multiple matrix regressions (partial Mantel tests, R package “phytools”) with 100,000 Mantel permutations to compute P-values. Phylogenetic relatedness, environmental distance and ancestral habitat difference were included as explanatory variables, and either D_tip_, D_max_, C_1_ or θ as response variables. The choice of variables to compare the magnitude of convergence across independent habitat transitions is extensively discussed in the supplementary information.

## Acknowledgments

We thank Camille Thomas-Bulle, Tanja Schwander, Anthony Lapsansky, William Toubiana and Guillaume Lavanchy for insightful discussions on the manuscript. This study would not have been possible without the work of many passionate phasmid enthusiasts and breeders who, over the years, documented the biology of many species. We are therefore very grateful to the amateur and professional phasmid community for publicly or privately sharing these notes and observations along with many high quality pictures. We are also thankful to Nicolas Cliquennois for insightful discussions regarding Malagasy and Mascarene stick insects. We finally thank Paul Bertner, Angus McNab, Chien C. Lee, Bruno Kneubühler, Nicolas Cliquennois and Paul Brock for permission to use their pictures.

This research was supported by NSF IOS-2015907 to DJE.

## Author Contributions

R.P.B., S.B. and D.J.E. designed research; S. B. curated and contributed specimens; R.P.B. performed the research; R.P.B. analyzed the data; R.P.B. and D.J.E. wrote the paper.

The authors declare that they have no conflicts of interest.

All data and code used in this study will be made available in a public repository (figshare).

## References

1. S. Conway Morris, The runes of evolution: how the universe became self-aware (Templeton Press, 2015).

2. G. R. McGhee, Convergent evolution: limited forms most beautiful (MIT Press, 2011) 10.4081/eb.2012.e2.

3. J. B. Losos, Convergence, adaptation, and constraint. Evolution (N Y) 65, 1827–1840 (2011).

4. D. B. Wake, M. H. Wake, C. D. Specht, Homoplasy: From detecting pattern to determining process and mechanism of evolution. Science (1979) 331, 1032–1035 (2011).

5. D. M. Grossnickle, et al., Incomplete convergence of gliding mammal skeletons. Evolution (N Y) 74, 2662–2680 (2020).

6. P. Trontelj, A. Blejec, C. Fišer, Ecomorphological convergence of cave communities. Evolution (N Y) 66, 3852–3865 (2012).

7. Š. Borko, P. Trontelj, O. Seehausen, A. Moškrič, C. Fišer, A subterranean adaptive radiation of amphipods in Europe. Nat Commun 12, 1–12 (2021).

8. R. G. Gillespie, S. P. Benjamin, M. S. Brewer, M. A. J. Rivera, G. K. Roderick, Repeated diversification of ecomorphs in hawaiian stick spiders. Current Biology 28, 941–947 (2018).

9. C. T. Stayton, The definition, recognition, and interpretation of convergent evolution, and two new measures for quantifying and assessing the significance of convergence. Evolution (N Y) 69, 2140–2153 (2015).

10. T. J. Ord, T. C. Summers, Repeated evolution and the impact of evolutionary history on adaptation. BMC Evol Biol 15, 1–12 (2015).

11. C. T. Stayton, Is convergence surprising? An examination of the frequency of convergence in simulated datasets. J Theor Biol 252, 1–14 (2008).

12. D. Esquerré, J. Scott Keogh, Parallel selective pressures drive convergent diversification of phenotypes in pythons and boas. Ecol Lett 19, 800–809 (2016).

13. Z. D. Blount, R. E. Lenski, J. B. Losos, Contingency and determinism in evolution: Replaying life’s tape. Science (1979) 362, eaam5979 (2018).

14. J. B. Losos, T. R. Jackman, A. Larson, K. De Queiroz, L. Rodríguez-Schettino, Contingency and determinism in replicated adaptive radiations of island lizards. Science (1979) 279, 2115–2118 (1998).

15. L. D. Mahler, T. Ingram, L. J. Revell, J. B. Losos, Exceptional Convergence on the Macroevolutionary Landscape in Island Lizard Radiations. Science (1979) 341, 292– 295 (2013).

16. P. F. Colosimo, et al., Widespread parallel evolution in sticklebacks by repeated fixation of ectodysplasin alleles. Science (1979) 307, 1928–1933 (2005).

17. S. J. Gould, Wonderful life: the Burgess Shale and the nature of history. (W. W. Norton & Company, 1989).

18. S. J. Gould, The structure of evolutionary theory. (Harvard University Press, 2002).

19. V. Orgogozo, Replaying the tape of life in the twenty-first century. Interface Focus 5, 20150057 (2015).

20. R. D. H. Barrett, D. Schluter, Adaptation from standing genetic variation. Trends Ecol Evol 23, 38–44 (2008).

21. G. L. Conte, M. E. Arnegard, C. L. Peichel, D. Schluter, The probability of genetic parallelism and convergence in natural populations. Proceedings of the Royal Society B: Biological Sciences 279, 5039–5047 (2012).

22. M. Bohutínská, et al., Genomic basis of parallel adaptation varies with divergence in Arabidopsis and its relatives. Proc Natl Acad Sci U S A 118, e2022713118 (2021).

23. S. Chaturvedi, et al., Climatic similarity and genomic background shape the extent of parallel adaptation in Timema stick insects. Nature Ecology & Evolution 2022 6:12 6, 1952–1964 (2022).

24. G. Montejo-Kovacevich, et al., Repeated genetic adaptation to altitude in two tropical butterflies. Nat Commun 13, 1–16 (2022).

25. I. S. Magalhaes, et al., Intercontinental genomic parallelism in multiple three-spined stickleback adaptive radiations. Nat Ecol Evol 5, 251–261 (2020).

26. R. D. H. Barrett, S. M. Rogers, D. Schluter, Natural selection on a major armor gene in threespine stickleback. Science (1979) 322, 255–7 (2008).

27. Y. E. Stuart, et al., Contrasting effects of environment and genetics generate a continuum of parallel evolution. Nat Ecol Evol 1, 1–7 (2017).

28. G. D. Ruxton, W. L. Allen, T. N. Sherratt, M. P. Speed, Avoiding attack: the evolutionary ecology of crypsis, aposematism, and mimicry. (Oxford University Press, 2019).

29. J. Skelhorn, H. M. Rowland, M. P. Speed, G. D. Ruxton, Masquerade: camouflage without crypsis. Science (1979) 327, 51 (2010).

30. G. O. Bedford, Biology and ecology of the Phasmatodea. Annu Rev Entomol 23, 125– 149 (1978).

31. S. Bradler, T. R. Buckley, “Biodiversity of Phasmatodea” in Insect Biodiversity: Science and Society, R. G. Foottit, P. H. Adler, Eds. (Wiley-Blackwell, 2018), pp. 281–313.

32. P. D. Brock, T. H. Büscher, Stick and Leaf-Insects of the World (NAP Editions, 2022).

33. T. R. Buckley, D. Attanayake, S. Bradler, Extreme convergence in stick insect evolution: phylogenetic placement of the Lord Howe Island tree lobster. Proceedings of the Royal Society B: Biological Sciences 276, 1055–1062 (2009).

34. S. Bradler, N. Cliquennois, T. R. Buckley, Single origin of the Mascarene stick insects: ancient radiation on sunken islands? BMC Evol Biol 15, 1–10 (2015).

35. S. Simon, et al., Old world and new world Phasmatodea: phylogenomics resolve the evolutionary history of stick and leaf insects. Front Ecol Evol 7, 1–14 (2019).

36. S. Bradler, J. A. Robertson, M. F. Whiting, A molecular phylogeny of Phasmatodea with emphasis on Necrosciinae, the most species-rich subfamily of stick insects. Syst Entomol 39, 205–222 (2014).

37. J. A. Robertson, S. Bradler, M. F. Whiting, Evolution of oviposition techniques in stick and leaf insects (Phasmatodea). Front Ecol Evol 6, 1–15 (2018).

38. N. Cliquennois, Révision des Anisacanthidae, famille endémique de phasmes de Madagascar (Phasmatodea: Bacilloidea). Annales de la Societe Entomologique de France 44, 59–85 (2008).

39. E. Tihelka, C. Cai, M. Giacomelli, D. Pisani, P. C. J. Donoghue, Integrated phylogenomic and fossil evidence of stick and leaf insects (Phasmatodea) reveal a Permian–Triassic co-origination with insectivores. R Soc Open Sci 7, 201689 (2020).

40. S. Bank, et al., Reconstructing the nonadaptive radiation of an ancient lineage of ground-dwelling stick insects (Phasmatodea: Heteropterygidae). Syst Entomol 46, 487– 507 (2021).

41. S. Bank, et al., A tree of leaves: phylogeny and historical biogeography of the leaf insects (Phasmatodea: Phylliidae). Commun Biol 4, 1–12 (2021).

42. S. Datta, S. Datta, Methods for evaluating clustering algorithms for gene expression data using a reference set of functional classes. BMC Bioinformatics 7, 1–9 (2006).

43. A. Camousseight, Revision taxonomica del genero Agathemera (Phasmatodea: Pseudophasmatidae) en Chile. Revista Chilena de Entomologia 22, 35–53 (1995).

44. C. Cubillos, A. Vera, Comparative morphology of the eggs from the eight species in the genus Agathemera Stål (Insecta: Phasmatodea), through phylogenetic comparative method approach. Zootaxa 4803, 523–543 (2020).

45. R. R. Junker, J. Kuppler, A. C. Bathke, M. L. Schreyer, W. Trutschnig, Dynamic range boxes – a robust nonparametric approach to quantify size and overlap of n-dimensional hypervolumes. Wiley Online Library 7, 1503–1513 (2016).

46. B. Blonder, et al., hypervolume: high dimensional geometry, set operations, projection, and inference using kernel density estimation, support vector machines, and convex hulls (2023).

47. J. Clavel, G. Escarguel, G. Merceron, mvmorph: an r package for fitting multivariate evolutionary models to morphometric data. Methods Ecol Evol 6, 1311–1319 (2015).

48. K. Arbuckle, C. M. Bennett, M. P. Speed, A simple measure of the strength of convergent evolution. Methods Ecol Evol 5, 685–693 (2014).

49. K. Arbuckle, A. Minter, Windex: Analyzing convergent evolution using the wheatsheaf index in R. Evolutionary Bioinformatics 2015, 11–14 (2015).

50. D. M. Grossnickle, et al., Challenges and advances in methods for measuring phenotypic convergence. bioRxiv, 2022.10.18.512739 (2023).

51. D. C. Collar, J. S. Reece, M. E. Alfaro, P. C. Wainwright, R. S. Mehta, Imperfect morphological convergence: Variable changes in cranial structures underlie transitions to durophagy in moray eels. American Naturalist 183 (2014).

52. M. Lässig, V. Mustonen, A. M. Walczak, Predicting evolution. Nat Ecol Evol 1, 1–9 (2017).

53. M. T. Wortel, et al., Towards evolutionary predictions: Current promises and challenges. Evol Appl 16, 3–21 (2023).

54. T. Leinonen, R. J. S. McCairns, G. Herczeg, J. Merilä, Multiple evolutionary pathways to decreased lateral plate coverage in freshwater threespine sticklebacks. Evolution (N Y) 66, 3866–3875 (2012).

55. M. Muschick, A. Indermaur, W. Salzburger, Convergent Evolution within an Adaptive Radiation of Cichlid Fishes. Current Biology 22, 2362–2368 (2012).

56. E. B. Rosenblum, C. E. Parent, E. E. Brandt, The molecular basis of phenotypic convergence. Annu Rev Ecol Evol Syst 45, 203–229 (2014).

57. J. F. Storz, Causes of molecular convergence and parallelism in protein evolution. Nat Rev Genet 17, 239–250 (2016).

58. A. J. Verster, A. K. Ramani, S. J. McKay, A. G. Fraser, Comparative RNAi screens in C. elegans and C. briggsae reveal the impact of developmental system drift on gene function. PLoS Genet 10, e1004077 (2014).

59. Y. M. Pacheco, “Ecomorph Convergence in Stick Insects (Phasmatodea) with Emphasis on the Lonchodinae of Papua New Guinea,” Brigham Young University. (2018).

60. R. Bouckaert, et al., BEAST 2: A Software Platform for Bayesian Evolutionary Analysis. PLoS Comput Biol 10, e1003537 (2014).

61. M. Chavent, V. Kuentz-Simonet, A. Labenne, J. Saracco, Multivariate Analysis of Mixed Data: The R Package PCAmixdata. ArXiv (2017).

62. G. Brock, V. Pihur, S. Datta, S. Datta, clValid: An R Package for Cluster Validation. J Stat Softw 25, 1–22 (2008).

63. A. Liaw, M. Wiener, Classification and Regression by randomForest. R News 2, 18–22 (2002).

64. L. J. Revell, phytools: An R package for phylogenetic comparative biology (and other things). Methods Ecol Evol 3, 217–223 (2012).

