## Supplementary information for "Divergence time and environmental similarity predict the strength of morphological convergence in stick and leaf insects"

**This PDF file includes:**

Supporting text
Figures S1 to S14
Tables S1 to S9
Legend for Movie S1
Legend for Dataset S1
SI References

**Other supporting materials for this manuscript include the following:**

Movie S1
Dataset S1

**Supporting Information Text**

**Supplementary Materials and Methods**

We used R software (v4.2.3) (1) for most analyses and data manipulation, as described below.

**DNA sequence alignment.** DNA sequences were obtained for 314 species of Phasmatodea and one species of Embioptera (the sister order of Phasmatodea (2)). We sampled regions of 7 genes: nuclear 18S rRNA (18S), 28S rRNA (28S) and histone subunit 3 (H3), and mitochondrial 12S rRNA (12S), 16S rRNA (16S), cytochrome-c oxidase subunit I (COI) and cytochrome-c oxidase subunit II (COII). Our species list was largely similar to the taxon list of a previous study (3) (Dataset S1). Sequences were extracted from Genbank and corresponded to data generated in previously published studies (3–8). They were reoriented and aligned using the R package "DECIPHER" (9). Protein coding sequences (i.e., H3, COI, COII) were first converted into amino acid sequences, aligned and reverse translated (*AlignTranslation*: "DECIPHER"; *R function*: "R package"). Frameshift errors occurring after the accidental insertion or deletion of one or two nucleotides during DNA sequencing were corrected after comparison of each sequence with an arbitrarily chosen unshifted sequence (*CorrectFrameshifts*: "DECIPHER"). Ribosomal genes (i.e., 18S, 28S, 12S and 16S) were first transcribed into RNA and then aligned while considering the secondary structure of non-coding RNA (*AlignSeqs*: "DECIPHER"). Gappy columns were then removed if they contained less than 5% nucleotide (*del.colgapsonly*: "ape")(10). As alignment extremities varied considerably in length, we trimmed each multiple sequence alignment (MSA) from the beginning and end using MACSE 2.05 (11) until a 30% nucleotide coverage was achieved (java -jar macse.jar -prog trimAlignment -align alignment.fasta -min_percent_NT_at_ends 0.3). All alignments were visually checked for conspicuously misaligned sections using Geneious v.2021.1 (www.geneious.com) and these were manually corrected if possible or otherwise excluded (n=13 excluded), and the MSAs were concatenated (*concatenate*: "apex")(12).

**Partition scheme.** To find the best-fitting partition scheme, we used PartitionFinder 2 (13) which separated the data into five subsets: mitochondrial rRNA 12S+16S (GTR+I+G+X), nuclear rRNA 18S+28S (GTR+I+G+X), protein codon positions 1+2 (GTR+I+G+X), COX position 3 (GTR+G+X), and H3 position 3 (HKY+I+G+X).

**Phylogenetic reconstruction.** Phylogenetic reconstructions were performed using this combined and partitioned dataset of 6,778 bp on the Cipres Science Gateway (www.phylo.org/)(14). Tree and divergence time estimation through Bayesian analyses were simultaneously performed in BEAST 2 (v. 2.6.3) (15). The optimal substitution model for each partition was selected using the bModelTest package in BEAST (v.1.2.1, "allreversible" (16)) and the partition scheme inferred by PartitionFinder.

We used 6 unambiguous crown-group phasmid fossils as minimum calibration points that were assigned to specific nodes on the tree using synapomorphy-based anatomical evidence (Table S1). We modeled all fossil constraints as a lognormal prior distribution with the minimum age as offset, a log-mean of 1.0 and a log-SD of 1.0. Tree and clock parameters were linked across all partitions and the analysis used an uncorrelated log-normal relaxed molecular clock model (UCLD) with a Yule model of speciation prior. We constrained our phylogenetic inferences to adopt the same backbone topology between major subclades as the most recent transcriptomic study (17). The analysis was run for 200 million generations with parameters and trees sampled every 500 generations. Convergence was assessed in Tracer v.1.7.1 (18) (ESS>200) and a Maximum Clade Credibility (MCC) tree was built in TreeAnnotator v.2.6.6 after the removal of the first 10% of trees as burn in.

**Morphological data.** We measured 1359 adult female specimens from 212 species included in the phylogeny. Males are less morphologically variable than females and are not considered in this study, as sexual dimorphism is extremely strong and variable across stick insect species and will be the focus of a companion study (Boisseau *et al.*, *in prep*). We gathered high quality photographs of live or pinned specimens in dorsal and/or lateral view from our own collection at the University of Göttingen (Germany), field guides (19–23), the published literature, and various online databases (24) (Dataset S1). Pictures were only included if the body of the specimen was relatively straight and undamaged. We measured between 1 and 18 different individuals per species (mean = 5.5 individuals per species). To standardize scale on each photograph (which were sometimes unscaled), we measured each of the 23 continuous traits described in Fig. S1 relative to body length using ImageJ (v1.51)(25). We collected data on body length (excluding ovipositor and subgenital plate) from the literature in addition to directly measuring it on properly scaled photographs (Dataset S1). We then took the median of each relative measurement across all females of a given species and we multiplied it by the median of body length to obtain absolute measurements for each species. We checked the congruence between data obtained from specimens in our collection at the University of Göttingen and specimens obtained from all other sources (Fig. S13), and between data collected from pinned specimens and from photographs of live animals (Fig. S14). In all cases, our measurements were highly congruent across sources.

Finally, we qualitatively scored body texture and overall body coloration. We assigned a separate, binary score to the dorsal surface of the mesothorax and abdomen for texture (0 = smooth, 1= rough and/or spiny). We qualitatively scored the dominant body coloration of each species using pictures of live specimens in captivity or in the wild (wings folded), as “green” (i.e., from light to dark green), “brown” (i.e., from light to dark brown, orange and black) or “other” (i.e., aposematic/contrasting coloration). Most phasmids exhibit colorations ranging from light green to dark brown or even black, while a few purportedly aposematic taxa have evolved conspicuous and contrasting coloration patterns including red, yellow or white.

Because body coloration is often extremely plastic in phasmids, we duplicated taxa that exhibited several color morphs that fit in separate categories, as well as taxa exhibiting a mixed brown/green color pattern or with no unambiguously dominant color. We acknowledge that such a broad qualitative assessment of coloration by human observers is limited and substantially biased as predators may display a different vision system (e.g., tetrachromatic vision of birds). However, more accurate and objective quantifications (e.g., using spectrograms) were beyond the scope of this study as they would require calibrated pictures and/or live material for a large number of species, and this is not possible at present. We intentionally kept the color categories broad, and assigned any ambiguous taxon to multiple categories, to avoid misclassifications. The raw datasets are available in Dataset S1.

**The phasmid morphospace.** To build a multidimensional morphospace, we performed a Principal Component Analysis (PCA) including continuous and categorical data (R package "PCAmixdata", v3.1)(26, 27). The analysis was performed on $log_{10}$-transformed continuous trait values, except for wing length and wing area (see below). We included body volume and body area as body size measurements. Then, we included the rest of the quantitative traits after controlling for the effect of size by substituting original trait values with the residuals calculated from a phylogenetically-corrected linear regression against body volume (*pgls*: "caper"). In these regressions, branch length transformations were optimized using maximum likelihood (lambda='ML'). Because wing length and wing area included zeros for wingless species, we divided the original measurements by body length or body length squared respectively, to obtain measures of relative wing length and area. We mean-centered and scaled to unit variance the continuous variables. In total, we included 23 continuous and three categorical variables (Fig. S1-2).

PCA results can be misleading when the data considered have a phylogenetic structure (28). Phylogenetic correction is not available for the PCAmix procedure, so we used another method to assess the effect of phylogenetic bias on some of our estimates of habitat hypervolumes on the morphospace (see below in subsection "Overlap between habitat categories on the morphospace"). The coordinates from the PCAmix (including both quantitative and qualitative variables) and the corresponding standard PCA (including only quantitative variables) exhibited very similar phylogenetic signals (average lambda over the first 15 PCs = 0.84 and 0.82 respectively) which suggested that using a phylogenetic PCA (*phyl.pca*:"phytools", method="lambda")(29) on only the continuous variables should provide a satisfactory estimate of phylogenetic bias in our analyses.

**Habitat data.** Despite being charismatic and popular in museums, private collections, and as pets, most phasmids have not been studied in the wild, and the habitat use of many species is surprisingly poorly understood (30). Most phasmids are active at night, and very little is known about these behaviors. However, it is during the daytime while these insects are resting that they are exposed to visually hunting predators (including entomologists), and collection records sometimes include information about the locations of these animals at the time they are found. Phasmids typically adopt two kinds of resting postures while hiding during the day. They either rest on or hang from diverse substrates. These postures likely reflect the type of camouflage they rely on: hanging species expose their bodies and are consequently more likely to rely on masquerade (i.e., mimicking of objects irrelevant to predators) while species that rest on surfaces are more likely to rely on crypsis through background matching. Stick insects can be found in the foliage of plants, in the leaf litter, on trunks, on logs, under bark or in tall grass (23, 30–32). We broadly classified the habitat use of species based on the typical resting posture and substrate of adult females. We surveyed the literature and field guides for observations of where each species is typically found, and iNaturalist (https://www.inaturalist.org) for observations in the field (see Dataset S1). Pictures in the field were only considered if the specimen was observed resting (with the forelegs in alignment with the rest of the body). We defined five habitat use categories: resting on the ground or trunks (including the base of trunks, mossy logs, under bark, in the leaf litter), resting on branches and leaves, hanging from branches and leaves, hanging from grass and resting on palm leaves. Grass and palm leaves were considered separately as they constitute rather distinct habitats (i.e., very thin leaves and stems and very tough wide leaves respectively). It should be recognized that this classification is broad and consequently does not account for the full diversity of substrates and host plants that phasmids live on.

**Definition of ecomorphs.** We next used our multidimensional morphospace data to cluster species into distinct ecomorphs by running a hierarchical clustering algorithm (*hclust*: "stats") using the Ward's method to define ecomorphs based on overall proximity on the morphospace (defined by the first 15 PC axes). We defined the optimal number of clusters thanks to the computation of a Biological Homogeneity Index (BHI, *clValid*: "clValid", validation= "biological") which measured how homogeneous clusters are based on habitat use (33, 34). This measure adapted from gene expression profile analyses corresponds to the average proportion of taxa pairs with matched habitat uses that are statistically clustered together based on their morphology (33). Clusters were defined by a fixed height threshold on the clustering dendrogram. The optimal number of clusters was then chosen as the minimal number of clusters to maximize BHI (Fig. S6). This procedure defined 20 separate groups of morphologically and ecologically similar taxa (thereafter called "ecomorphs") (Fig. 4, Fig. S3-5).

To describe the morphological niche occupied by each ecomorph, we identified the morphospace axes that best distinguished them (Table S4). Using the full dataset, we trained random forest models to classify a species in either a given ecomorph of interest or in a different one, given the first ten axes of the morphospace (*randomForest*: "randomForest")(35). This machine learning method classifies species into either belonging to an ecomorph of interest or not while making minimal assumptions regarding the shape of the regions occupied by these two groups on the morphospace. It also accounts for interactions between trait axes. The algorithm uses decision trees that partition the morphospace into a set of non-overlapping rectangular hypervolumes inside which the group heterogeneity is minimized. Internal nodes in a decision tree correspond to a split along one dimension of the morphospace and terminal nodes correspond to a unique non-overlapping hypervolume. The group of each species is then "voted" by each decision tree in the random forest based on its coordinates on the morphospace. Species are then assigned to a group by counting the votes of the forest of decision trees. After training on the full dataset, the algorithm evaluates its classification accuracy by comparing its predictions relative to the observed group of each species. It then estimates the relative importance of each dimension to distinguish a given ecomorph by quantifying the decrease in accuracy followed by the omission of a given dimension. Thus, we could identify the axes of the morphospace that were relatively more important in distinguishing each ecomorph from the rest (Table S4).

**Ancestral state reconstruction of ecomorphs.** Ecomorphs, as defined by our hierarchical clustering analysis, were mapped on the MCC tree to establish whether they had single or multiple origins (Fig. 3). We ran ancestral state reconstructions using stochastic character mapping as implemented in the R package "phytools" (function *make.simmap*) (29). Given the large number of ecomorphs (n=20), only the "equal rate" transition model (assuming a single transition rate between ecomorphs) could be run. The transition matrix was calculated using MCMC, the prior distribution on the root node of the tree was estimated and 1,000 stochastic character maps were subsequently simulated and summarized to get posterior probabilities for each state at each node.

**Visualizing morphological convergence.** To visualize how some lineages evolved similar trait values, we reconstructed ancestral trait values at each node on the MCC tree (*fastAnc*, “phytools”)(29) and extrapolated trait values from node to node to estimate ancestral values along all the branches of the tree at 500,000 year time intervals (36). We built a dynamic 2-D phylomorphospace (including PC1 and PC2) to show how lineages diversified morphologically over time (Video S1). At each time step, the ecomorph of each lineage was predicted given its position on the morphospace using a random forest model, initially trained with the full extant dataset and the first five dimensions of the morphospace.

**Ancestral state reconstruction of habitat use.** Habitat use was mapped on the MCC tree to uncover the number of transitions toward each of the five categories (Fig. 5A). We ran ancestral state reconstructions using stochastic character mapping (*make.simmap*: “phytools”). The transition matrix was calculated using maximum likelihood and using an all-rates-different model (model= “ARD”). 1,000 stochastic character maps were subsequently simulated and summarized to get posterior probabilities for each state at each node.

**Overlap between habitat categories on the morphospace.** To quantify morphospace occupation by species exhibiting different habitat uses, we estimated 6-dimensional hypervolumes (PC1-PC6, accounting for 81% of the total variation) using dynamic range boxes (R package "dynRB")(37). This non-parametric quantile-based approach does not assume any particular distribution of the data but rather considers the distribution of the observed values and is robust to outliers (37). To calculate volumes, it calculates the range sizes of each habitat for each dimension and aggregates them by either calculating their product, arithmetic or geometric mean. The overlap between hypervolumes is calculated as the portion of the hypervolume of habitat A covered by the hypervolume of habitat B (port(A,B)) and vice versa (port(B,A)). Hypervolumes and overlap values are bounded between 0 and 1. Using heatmaps, we represented pairwise overlaps between habitats for the full hypervolumes and for the five dimensions individually, in order to visualize the morphospace axes best differentiating the five habitat categories (Fig S9). Estimated range size for each habitat was also plotted for the full niche and the 6 dimensions individually (Fig S7).

We also assessed pairwise similarity of the habitat hypervolumes reconstructed through high-dimensional kernel density estimations using other indexes implemented in the R package “hypervolume”(38). First, we computed the Jaccard similarity index which quantifies the ratio of the intersection to the union of the hypervolumes, the Sørensen–Dice similarity index which measures twice the size of their intersection relative to the sum of their individual sizes, and the volumes of shared components of the two hypervolumes divided by their respective size. Therefore, these indexes provide a normalized assessment of the overlap between two hypervolumes. We also quantified the Euclidean distance between the hypervolume centroids and the minimum Euclidean distance between two sets of random points comprising the two hypervolumes (Fig. S10).

To test whether we could predict the habitat of a species given its position on the morphospace, we used machine learning random forest models (R package "randomForest" v4.6-14)(39). The algorithm was first trained (number of trees=500, number of variables tried at each split =2, maximum number of terminal nodes = 50) on a randomly downsampled dataset including 75% of all species with available morphological and habitat data (n=274 taxa including duplicated species showing color polymorphism), and then used to predict habitat for the remaining 25% of the data. We compared the predictive performance of the models (i.e., classification accuracy) including one to 15 morphospace axes and repeated this step for 1,000 randomly sampled datasets for each set of the morphospace axes, and averaged prediction accuracies (Fig. 5C-D).

Given that the PCAmix procedure used to build the morphospace does not allow for phylogenetic correction, we assessed the effect of phylogenetic bias on volume estimates for each habitat following Nordén et al., 2019. Using the "hypervolume" package, we compared habitat hypervolumes (calculated from kernel density estimates using the first 15 PCs, function *hypervolume_box*) estimated by a standard PCA (*prcomp*: "stats") with those estimated using a phylogenetic PCA (*phyl.pca*: "phytools", method="lambda"), excluding the qualitative variables texture and color. Overall, volume estimates and patterns were very similar between the standard and phylogenetic PCAs (Fig. S8).

**Process-based tests of convergence – evolutionary model fitting.** To test for morphological convergence among lineages that independently transitioned to the same habitat, we fit multivariate models of continuous trait evolution to PC1-PC5 (77% of the total variance) using the "mvMORPH" R package (41). "mvMORPH" takes the correlation between PC-scores into account when fitting models. Goodness-of-fit of the models was evaluated using the small-sample corrected Akaike information criterion (AICc). Log-likelihood was computed using the "sparse" method during model-fitting.

We first fitted the single-regime Brownian motion (BM1), Ornstein–Uhlenbeck (OU1), and early burst (EB) models (fucntions *mvBM*, *mvOU* and *mvEB* respectively) which represent the null hypothesis. Support for these models would indicate that habitat use does not strongly affect phasmid morphology. The BM1 model includes a single phylogenetic mean and evolutionary rate ($\sigma^2$) to model stochastic trait evolution. The OU1 model considers the action of selection by including a parameter, α, that corresponds to the strength of attraction toward a trait optimum (θ), in addition to $\sigma^2$. The EB model uses a decay parameter to model a decrease in evolutionary rate with time.

We tested for the presence of distinct evolutionary regimes for different habitats using multi-regime models, which allow the model parameters to vary between groups. For each habitat category, we fitted 2-regime models where the given habitat category was considered its own regime while the rest belonged to the other regime. We also ran 5-regime models which considered each habitat as a separate evolutionary regime. The ancestral histories for each of the tested regime assignments were reconstructed on the MCC tree using 100 stochastic character maps. These maps were generated with the *make.simmap* function in the "phytools" R package (29). The transition matrix was calculated using maximum likelihood and using an all-rates-different model (model= "ARD"). We fitted the evolutionary models to the 100 "simmap" trees and averaged the results. Two types of multiple-regime BM models were fit. First, we allowed phylogenetic means to vary among regimes but kept $\sigma^2$ constant (BM1m). Second, we allowed both phylogenetic means and evolutionary rates to vary among regimes (BMMm). Multiple-regime OU models (OUM) allowed trait optima to vary among regimes, but $\sigma^2$ and $\alpha$ remained constant (Table S5). Support for OUM models would suggest selection for different trait optima corresponding to different habitats, possibly due to unique adaptive peaks. In contrast, support for BM1m or BMMm models would suggest that different habitat categories correspond to separate morphospatial zones but not to single adaptive peaks.

Finally, we fitted 20-regimes BM1m, BMMm and OUM models with regimes corresponding to ecomorphs (Table S6). Support for the OUM model would suggest that ecomorphs correspond to unique adaptive peaks.

**Pattern-based tests of convergence.** In parallel to the process-based tests of convergence described above, we also performed pattern-based tests and quantifications of convergence (Table S7). For each habitat use category, we first calculated the Wheatsheaf index ($w$), corresponding to the ratio of the average pairwise phenotypic distance between all taxa in the tree, to the average pairwise distance between all putatively convergent taxa, correcting for phylogenetic relatedness (42). We used the R package "windex" to calculate $w$ for each morphotype (43). The first five morphospace axes (PC1-PC5) were used. For each habitat category, we compared the obtained index to the ones calculated from random distributions of phenotypic values on the same tree topology obtained from 1,000 bootstrap resamples. The associated p-values test the hypothesis that convergence is significantly stronger for that habitat category than would be expected by chance.

We then calculated four convergence metrics, C1-C4, using the "convevol" R package (44) for each habitat category. In contrast with $w$, C1-C4 measure the increase in similarity between the convergent taxa through time and are therefore useful to distinguish convergent evolution from stasis, which the Wheatsheaf index is incapable of doing. C1-C4 use ancestral state reconstruction via a Brownian model of trait evolution to compare the distances between phylogenetic tips in the phylomorphospace and the distances between ancestral nodes. C1 is calculated as $C1 = 1 - \frac{D_{tip}}{D_{max}}$, where $D_{tip}$ is the present Euclidean distance between a given pair of taxa of interest on the morphospace and $D_{max}$ the maximum distance between any two pairs of taxa along those two lineages (extant or ancestors) (Fig. 6C). C1 therefore ranges from 0 to 1 and quantifies the phenotypic distance that has been reduced by convergent evolution over time. C1 close to 1 indicates strong convergence as it suggests that the extant taxa are much more phenotypically similar than were their ancestors. Alternatively, it can suggest that convergent lineages moved a relatively large distance on the morphospace in a similar direction relative to the present distance between these lineages (45). C2, calculated as $C2 = D_{max} - D_{tip}$, measures the absolute magnitude of convergence. C3 and C4 are standardized versions of C2. C3 measures the magnitude of convergence relative to the sum of phenotypic distances travelled on the morphospace by all the lineages starting from the common ancestor of the two taxa of interest, and is calculated as $C3 = \frac{C2}{L_{tot,clade}}$. And finally, $C4 = \frac{C2}{L_{tot,tree}}$ quantifies the amount of convergence scaled by the total phenotypic evolution in the entire tree. It should be noted that C1-C4 can only be calculated for pairs of taxa. Therefore, for each habitat category, we calculated and averaged C1-C4 for all possible pairs of taxa corresponding to separate independent transitions toward the given habitat category. To assess the significance of obtained C1-C4 values, we ran 1,000 simulations of character evolution along the phylogeny using a Brownian Motion model with the variance–covariance matrix determined from the data. Returned p-values test the hypothesis that convergence is stronger than would be expected randomly (i.e., with no constraint on the direction of evolution).

C1-C4 may be high when convergent lineages diverged substantially after splitting and subsequently re-evolved similarities, or when convergent lineages shifted in parallel towards the same direction on the morphospace (45). To distinguish those two scenarios, we computed the recently developed $C_t$ measures, which compare the extant phenotypic distance between the convergent lineages ($D_{tip}$) to the maximum reconstructed ancestral distance at a given time point ($D_{max,t}$) during their evolution (i.e., between synchronous points along the evolutionary trajectories)(45). Unlike C1-C4, Ct1-Ct4 are only expected to be high when lineages moved away from one another at some point in their evolutionary history and subsequently got closer. We calculated Ct1-Ct4 and associated p-values in a similar fashion as for C1-C4 using the function *convSigCt* in "convevol".

Finally, we quantified parallelism in the evolutionary trajectories of convergent lineages by calculating the angle ($\theta$) between these trajectories on the morphospace (Fig. 6C). We reconstructed the trajectories of convergent lineages from the position of the node immediately prior to the inferred habitat transition, to that of the tip of interest. Pairwise angles were calculated using the function *angleTest* in the R package "Morpho" (46). The procedure to calculate $\theta$ and associated p-values were similar to the ones used for C1-C4 and Ct1-Ct4. In this case, p-values test the hypothesis that $\theta$ is lower (i.e., trajectories are more parallel) than would be expected by chance.

**Environmental data.** We gathered information about the geographic range of each species based on sampling location of type specimens and observations on iNaturalist (available from https://www.inaturalist.org, accessed July 2021). For each species, we then selected the median location with the most central latitude. From the GPS coordinates of the most central location for each species, we extracted data on annual mean temperature, mean diurnal range (i.e., mean of monthly (maximum - minimum temperature)), temperature seasonality (i.e., standard deviation ×100), maximum temperature of warmest month, minimum temperature of coldest month, annual temperature range (i.e., maximum temperature of warmest month - minimum temperature of coldest month), annual precipitation, precipitation of wettest month, precipitation of driest month, precipitation seasonality (i.e., coefficient of variation) from worldclim (47) (available from https://www.worldclim.org/, Accessed July 2021). We also extracted the length of the growing period (i.e., number of days during a year when temperatures are above 5°C and precipitation exceeds half the potential evapotranspiration (48), available from https://data.apps.fao.org/map/catalog), the total annual growing degree days (i.e., a measure of the annual amount of thermal energy available for plant and insect growth; Climate Research Unit, Univ. of East Anglia, available from https://sage.nelson.wisc.edu/data-and-models), and the Net Primary Production of biomass (NPP) (grams of dry matter per $m^2$ per year; Climate Research Unit,

Univ. of East Anglia, period 1976-2000, available from https://data.apps.fao.org/map/catalog) from the FAO Map Catalog. NPP was used as a proxy for vegetation density and overall food availability. In parallel, we extracted bird, mammal and amphibian species richness at the central location of each species range from the IUCN (International Union for Conservation of Nature) database (available from https://www.iucnredlist.org/, accessed in February 2023). These were used as a proxy for predator diversity. Finally, we scored the vegetation layer in which each species is typically found during the day by surveying the scientific literature, field guides and online databases (see the habitat data section). We defined three broad ordinal categories: ground/shrub [1], shrub/understory [2] and understory/canopy [3]. These vegetation layers included overlapping names to emphasize that boundaries between inhabited vegetation layers are loose as phasmids often navigate between them. Ground/shrub dwelling taxa were defined as often resting below 1.5 meters above the ground. Understory/canopy dwelling taxa are typically found high up in the tree canopies (>5 meters above the ground) and rarely close to the ground. Finally, shrub/understory species are typically found at intermediate heights in shrubs, in the canopy of small trees, or on the trunk of large trees but do not climb higher than four or five meters. We summarized the variation in the 17 environmental variables by running a principal component analysis (*prcomp*, “stats”)(Fig. S11).

**Hypervolumes of habitat categories on the environmental space.** To quantify how variable environmental conditions are between species occupying the same habitat category, we estimated the hypervolume size of each habitat category on the environmental space (Fig. S12). We estimated 6-dimensional hypervolumes (PC1-PC6, accounting for 91% of the total environmental variation) using dynamic range boxes (aggregation method = product, R package "dynRB")(37) and Kernel density estimations (R package "hypervolume") (38).

**Analysis of the variation in the extent of morphological convergence.** We tested the effects of three factors on the strength of morphological convergence: the phylogenetic relatedness between the convergent lineages, their environmental distance, and whether they started from the same ancestral habitat. We only considered the repeated transitions toward resting on the leaf litter and trunks (n=15, Fig. 6A) and towards resting on branches and leaves (n=14, Fig. 6B) for these analyses. Transitions towards other habitat categories were too rare to allow sufficient statistical power ($n \leq 4$, Fig. 5A). For both habitat categories separately, phylogenetic relatedness, environmental distance, ancestral habitat difference and morphological convergence were computed for all possible pairs of taxa corresponding to separate independent transitions toward the habitat category, and then assembled as distance matrices. Pairwise phylogenetic relatedness was estimated as the age of the most recent common ancestor of the two lineages. Pairwise environmental distance was calculated as the Euclidean distance on the environmental PC1-PC4 (accounting for 80.1% of the total environmental variation). Pairwise ancestral habitat difference was scored as either 0 if both lineages transitioned to the habitat of interest from the same ancestral habitat, or 1 otherwise. Finally, to quantify morphological convergence we computed pairwise $D_{tip}$, pairwise $D_{max}$, pairwise $C_1$ and pairwise $\theta$.

We fitted multiple matrix regressions (partial Mantel tests, *multi.mantel*, “phytools”) with 100,000 Mantel permutations to compute P-values. Phylogenetic relatedness, environmental distance and ancestral habitat difference were included as explanatory variables, and either $D_{tip}$, $D_{max}$, $C_1$ or $\theta$ as response variables.

**Definitions and choice of convergence metrics.** In this study, we adopt a rather broad definition of convergence: the independent evolution of similar phenotypes in multiple lineages. Therefore, this definition includes lineages that evolved to be more similar to one another than their ancestors were to each other (convergence *sensu* (44, 49, 50); and lineages that shifted in the same direction in the phenotypic space (i.e., parallel shifts (51)), sometimes ending up less similar to one another than their ancestors were (i.e., “imperfect” convergence (52)). Quantitative tests for phenotypic convergence have multiplied with the recent advent of novel methods (42, 44, 53, 54). The C-measures are popular metrics for identifying and quantifying morphological convergence (44). C-measures compare the distance between focal phylogenetic tips ($D_{tip}$) on the morphospace and the maximum distance between the focal lineages at any points along their reconstructed evolutionary trajectories ($D_{max}$) (Fig. 6C). While originally designed to specifically identify cases where lineages evolve to be closer than their ancestors were, C-measures actually fail at differentiating these cases from parallel shifts and imperfect convergence (45). C-measures increase when $D_{tip}$ decreases (i.e., phylogenetic tips are closer on the morphospace) and when $D_{max}$ increases. $D_{max}$ will be high when lineages diverged substantially on the morphospace at one point during their history before coming back to the same region. But $D_{max}$ will also be high when lineages move across long distances on the morphospace in a similar direction. Therefore, high C-measures indicate that lineages are currently close on the morphospace relative to either their maximal ancestral distance, or the distance they travelled on the morphospace after splitting. This is why we argue that C-measures are appropriate metrics of the extent or strength of morphological convergence, considering our broad definition. $C_t$-measures were recently developed to specifically differentiate between these scenarios by restricting $D_{max}$ to synchronous ancestral nodes along the evolutionary trajectories of the focal lineages (45). By doing so $D_{max}$ will only be relatively high if, at some point in their history, ancestors of focal lineages were relatively far apart on the morphospace. Application of this method to stick insect morphological evolution gave very low values (Table S7) indicating that most of the convergence associated with similar habitat transitions happened through parallel evolutionary changes. This was further confirmed by measurements of the angle between the convergent trajectories that showed more parallel trajectories than expected by chance (Table S7).

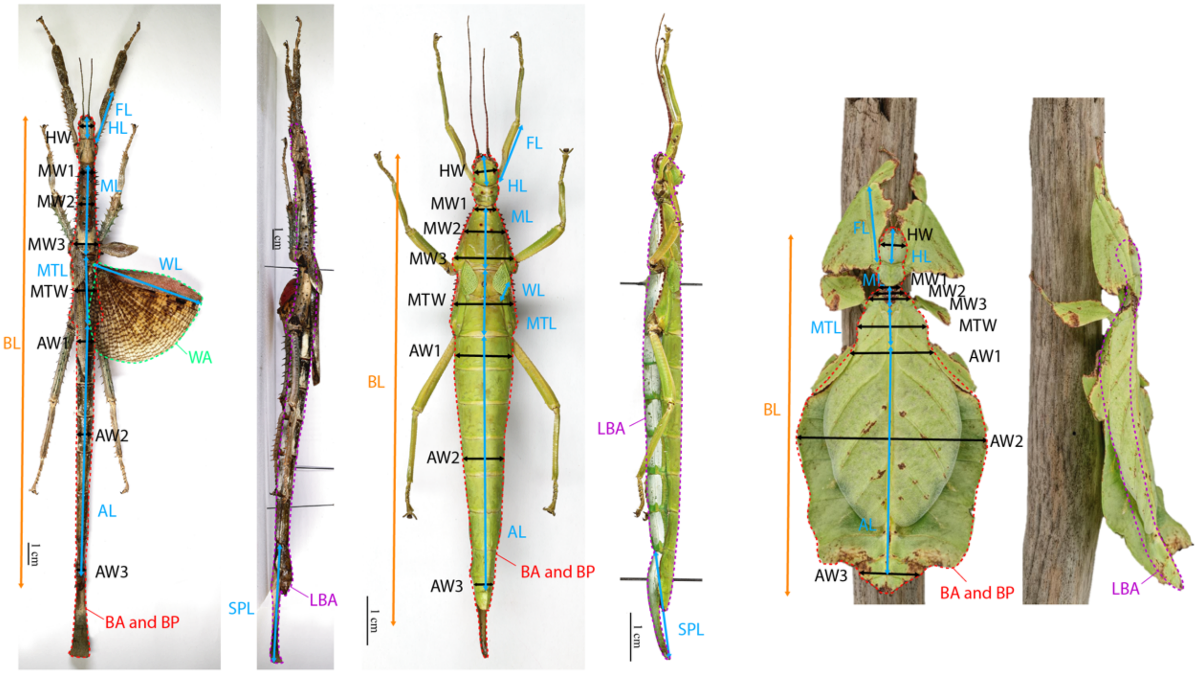


**Whole body measurements:**

BL: Body length (mm)
BA: Body dorsal area (mm²)
BP*: Body dorsal perimeter (mm)
LBA: Body lateral area (mm²)

BC: Body circularity $= 4\pi \times \frac{BA}{BP^2}$

BS: Body Solidity $= \frac{BA}{\text{Convex hull area*}}$

BW: Body Width (mm)$= \frac{BA}{BL^2}$

BH: Body Height (mm)$= \frac{LBA}{BL^2}$

BV: Body Volume (mm³) $= \frac{1}{4} \times \pi \times BW \times BH \times BL^3$

**Length (mm):**

FL: Front femur length
HL: Head length
ML: Mesothorax length
MTL: Metathorax length
AL: Abdomen length (from 2nd to 9th segment)
SPL: Subgenital plate length

**Width (mm):**

HW: Head width
MW1: Mesothorax width front
MW2: Mesothorax width middle
MW3: Mesothorax width rear
MTW: Metathorax width middle
AW1: Width 2nd abdominal segment
AW2: Width 5th abdominal segment
AW3: Width 9th abdominal segment

**Wing:**

WL: Hindwing length (mm)
WA: Hindwing area (mm²)

**Figure S1** : Quantitative morphological measurements of phasmid specimens. **Left:** Adult female *Achrioptera punctipes cliquennoisi* Hennemann & Conle 2004 (Specimen MNHN-EO-PHAS127, project RECOLNAT (ANR-11-INBS-0004), photographs by Marion Depraetere, 2015, CC-BY-NC-ND). **Middle:** Adult female *Diapherodes martinicensis* Lelong & Langlois, 2005 (Specimen MNHN-EO-PHAS542, project RECOLNAT (ANR-11-INBS-0004), photographs by Marion Depraetere, 2015, CC-BY-NC-ND). **Right:** Adult female *Pulchriphyllium giganteum* Hausleithner, 1984 (Culture “Tapah Hills”, Bruno Kneubuehler, Switzerland, photographs by Bruno Kneubuehler, used with permission).

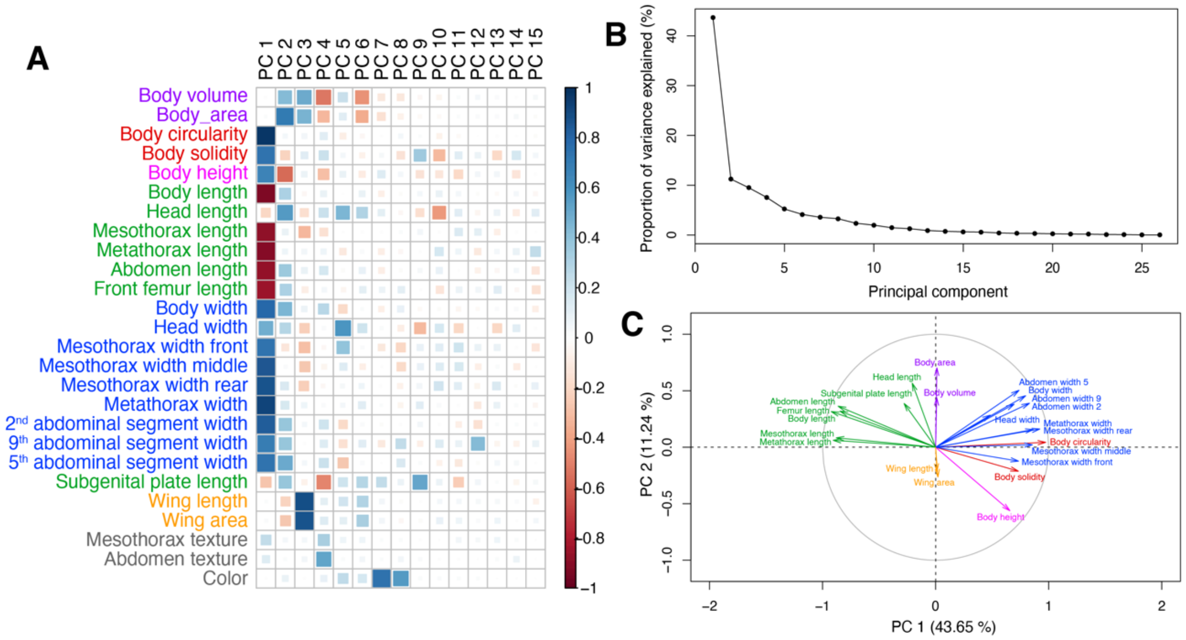


**Figure S2** : Results of the PCAmix analysis of 24 quantitative measurements and three qualitative scores of body mesothorax and abdomen texture, and overall body coloration. **A**: correlation plot between the 15 first principal components and all the original dataset features. **B**: Proportion of total variance explained by each principal component. **C**: Loading plot for the first principal components and the quantitative dataset features.

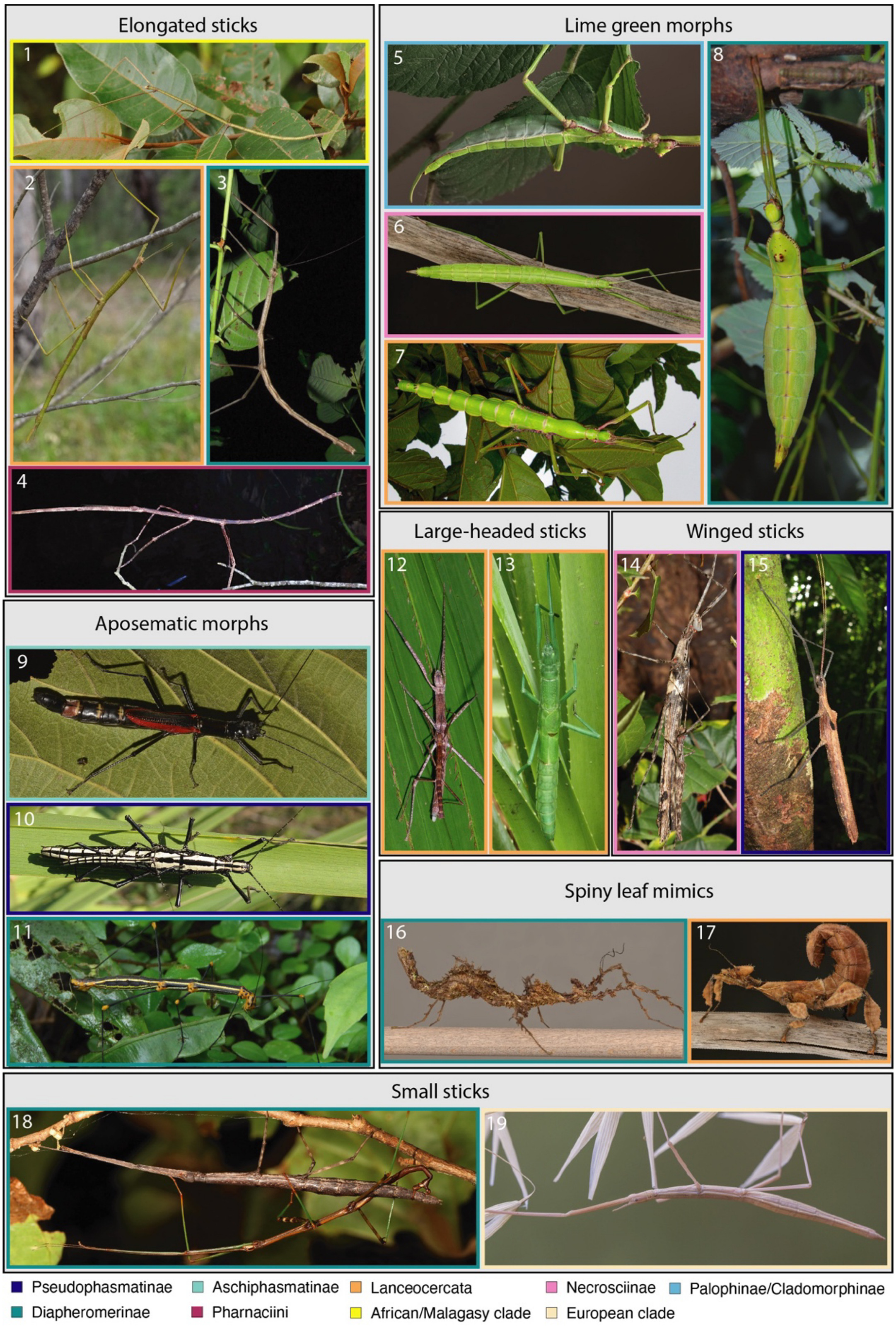


**Figure S3**: Photographs of adult females of each ecomorph (part 1/3). The color surrounding each picture corresponds to a major phylogenetic clade. The list of species and photograph information can be found in Table S9.

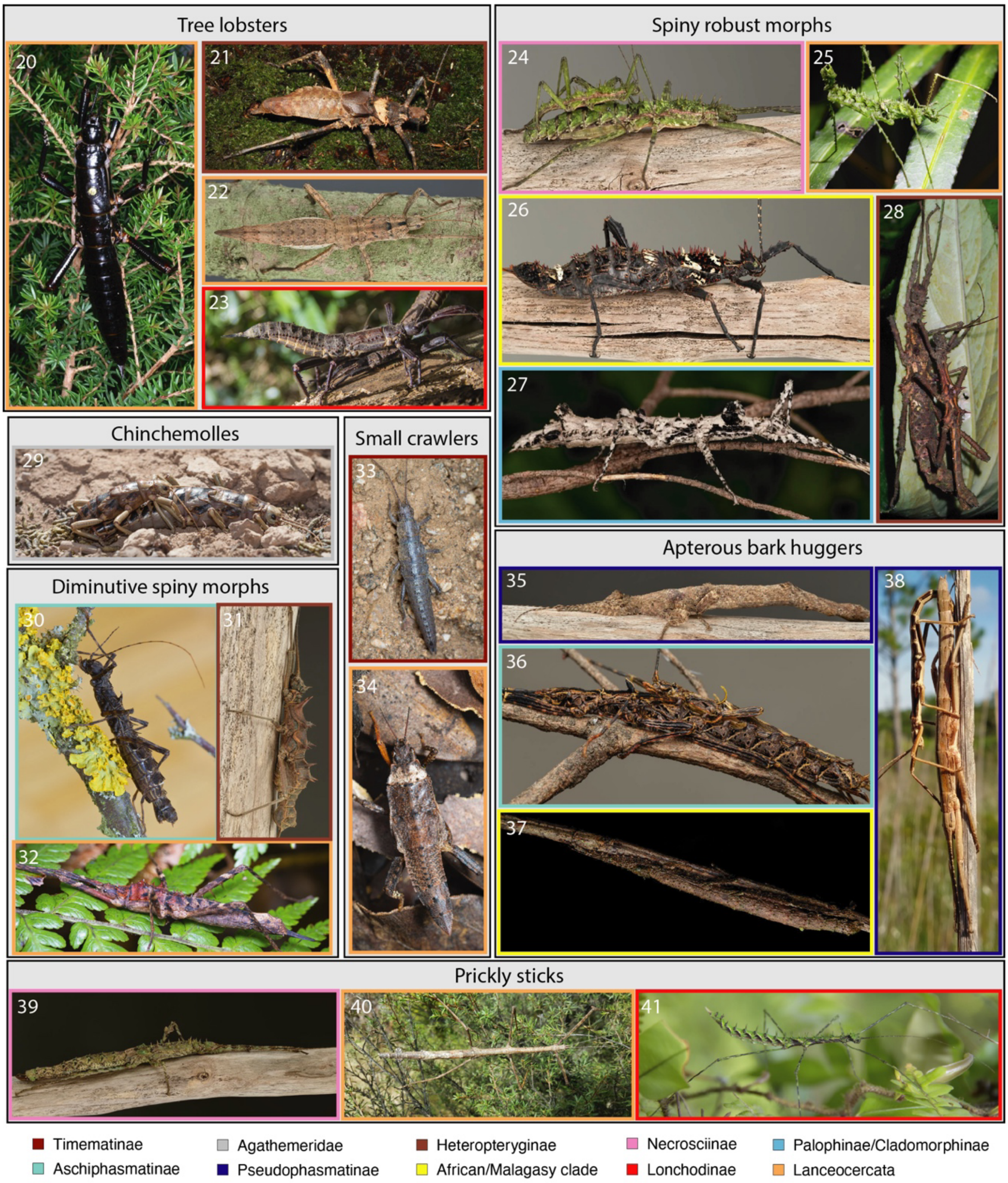


**Figure S4**: Photographs of adult females of each ecomorph (part 2/3). The color surrounding each picture correspond to a major phylogenetic clade. The list of species and photograph information can be found in Table S9.

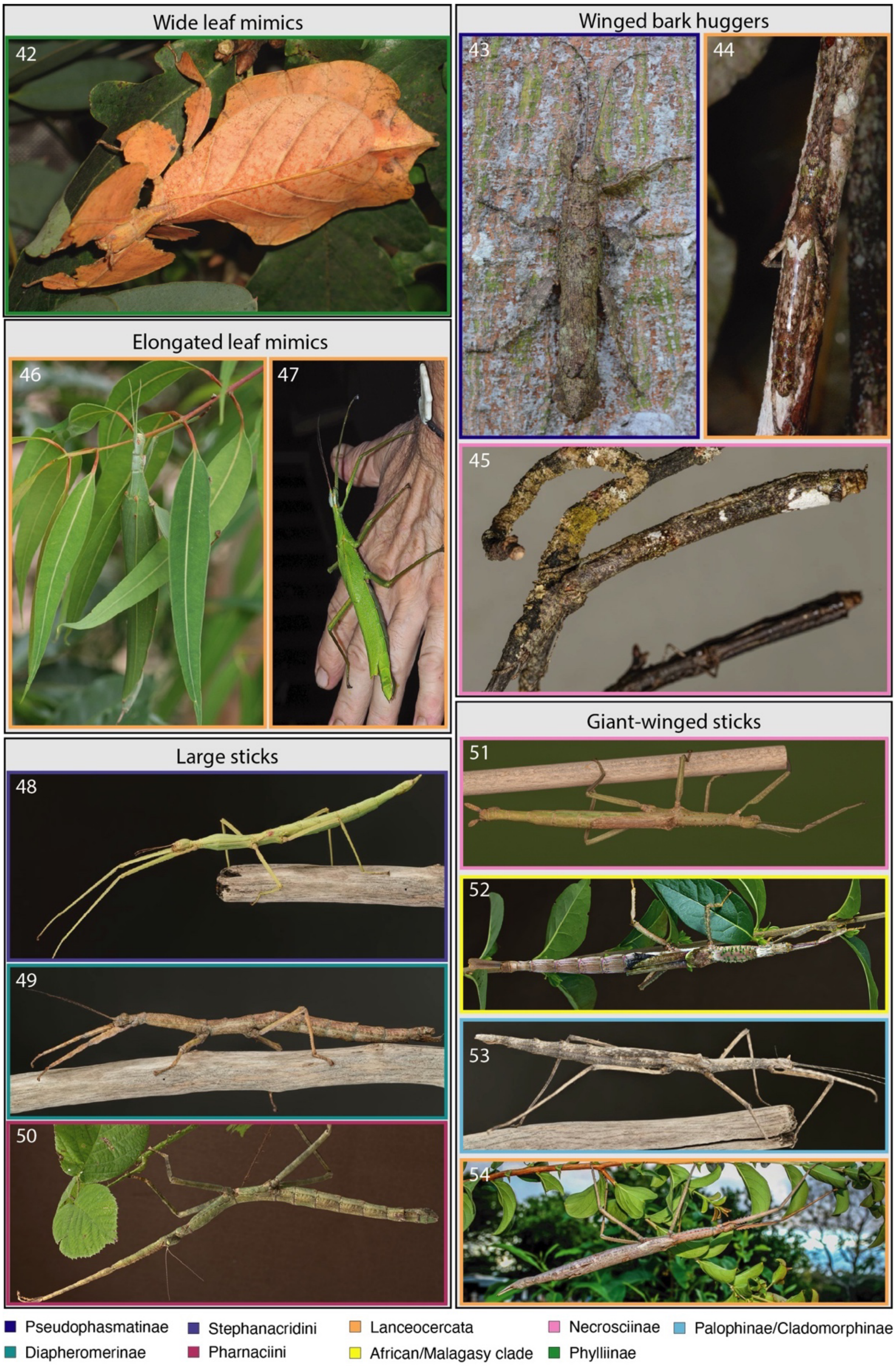


**Figure S5**: Photographs of adult females of each morphotype (part 3/3). The color surrounding each picture correspond to a major phylogenetic clade. The list of species and photograph information can be found in Table S9.

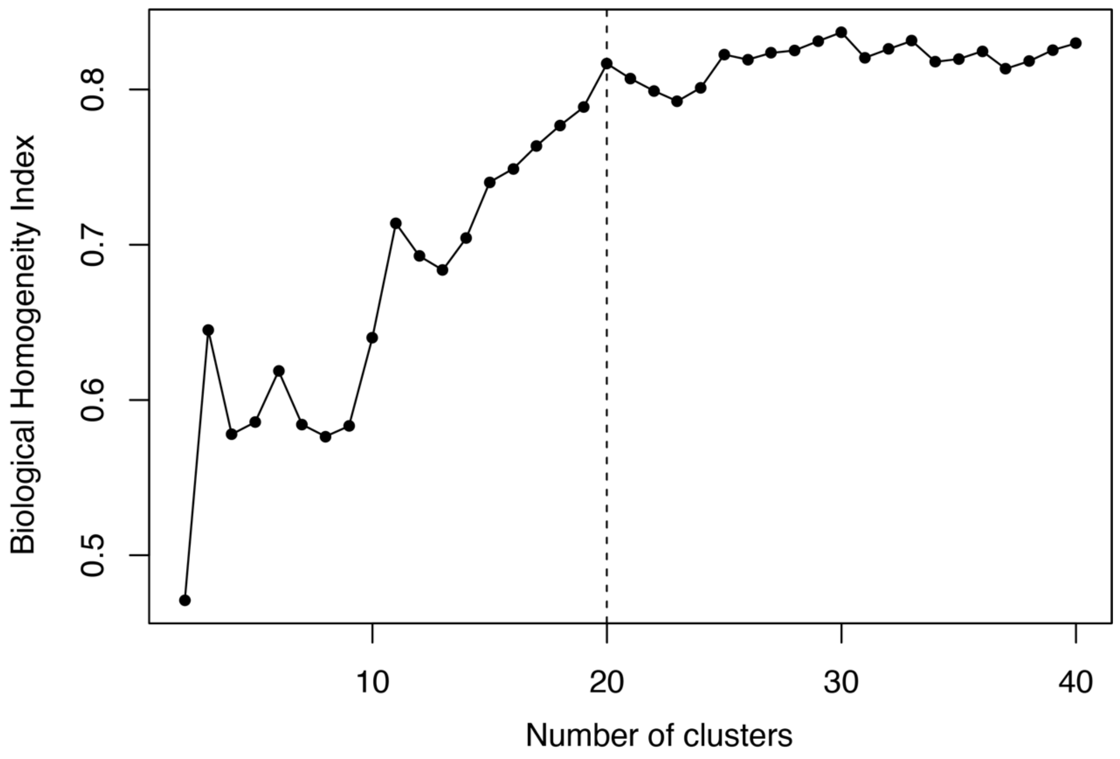


**Figure S6**: Biological homogeneity index (BHI, as calculated by R package "clValid") as a function of the number of cluster defined from the morphological hierarchical clustering dendrogram (Fig. 4). The BHI is calculated as the average proportion of taxa pairs with matched habitat uses that are statistically clustered together based on their morphology (i.e., proxy for habitat use homogeneity within morphological clusters). The optimal number of clusters was chosen as the minimal number of clusters to maximize BHI and is represented by the dashed line.

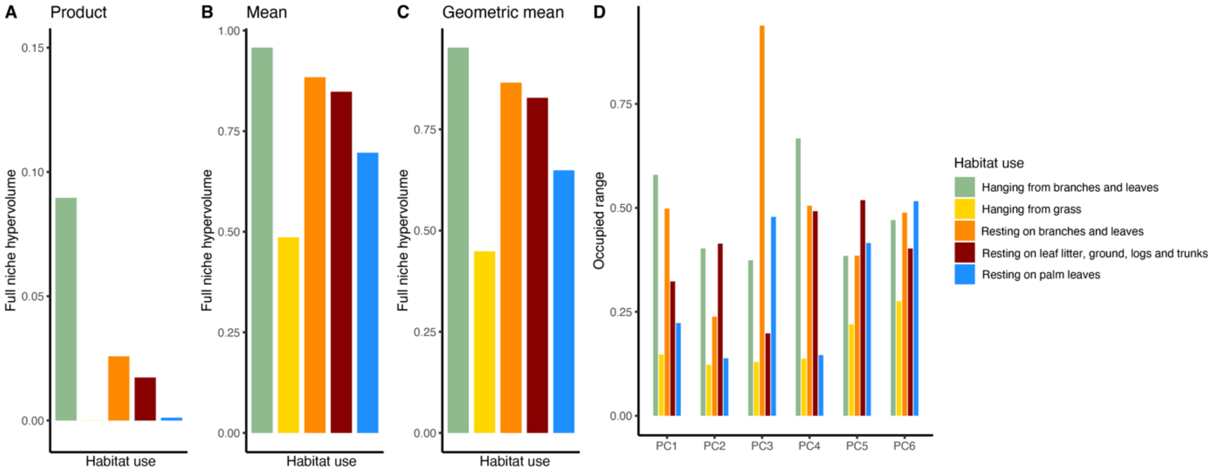


**Figure S7**: Morphospatial hypervolume occupied by each habitat categories as estimated using dynamic range boxes (R package “dynRB”). **A-C**: 6-dimensional hypervolume size for each habitat category obtained using three different aggregation methods: product (**A**), arithmetic mean (**B**) and geometric mean (**C**). **D:** Occupied range size by each habitat category along the first 6 axes of the morphospace.

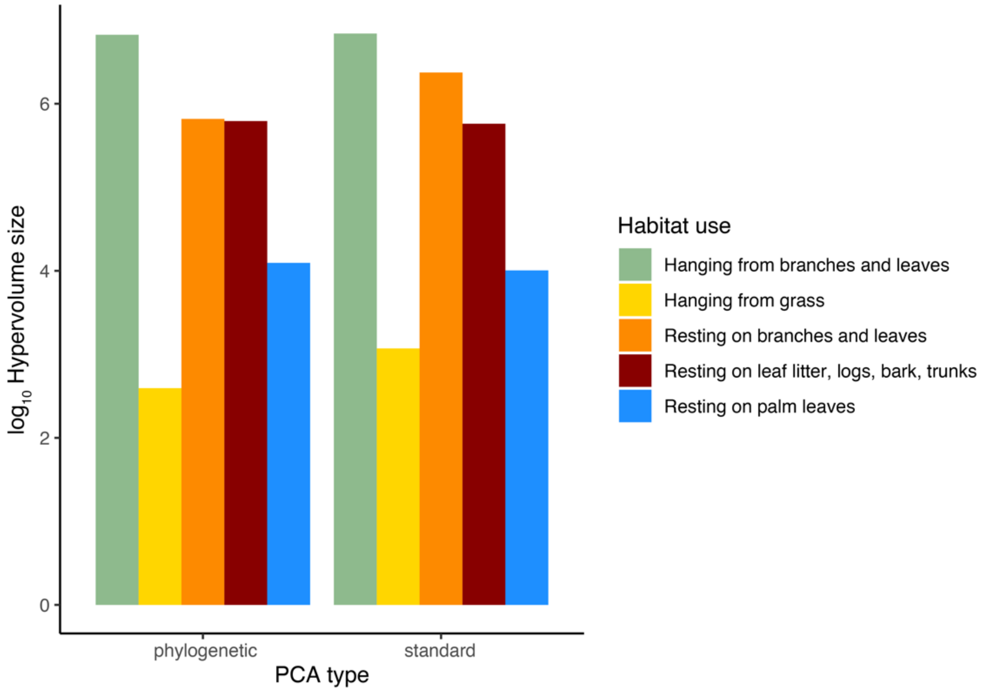


**Figure S8**: Habitat hypervolume size on the morphospace, estimated from kernel density estimates using the first 15 principal component scores derived from a standard PCA or a phylogenetic PCA.

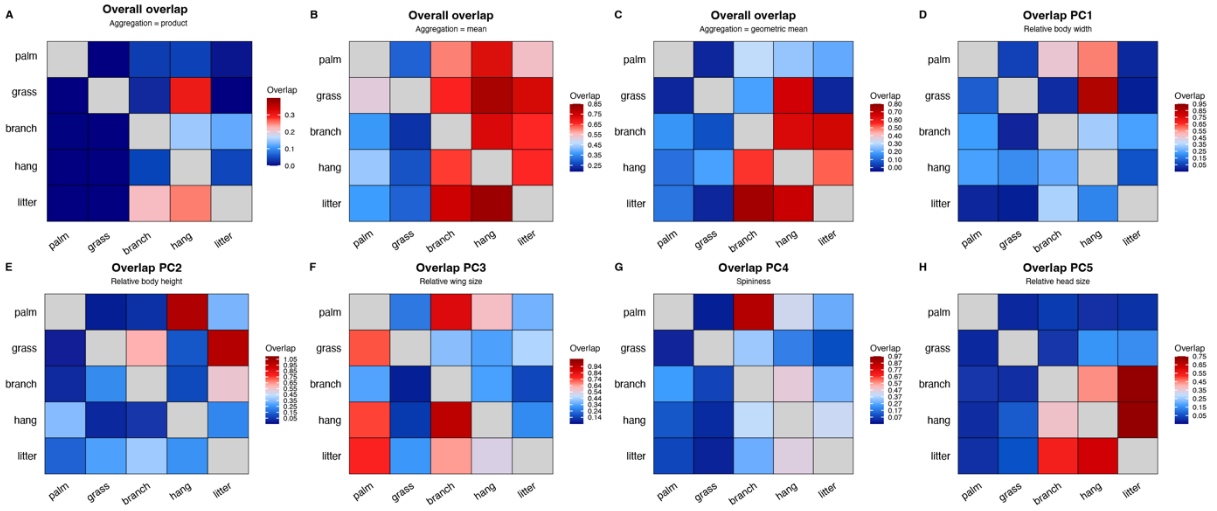


**Figure S9**: Morphospace overlap between different habitat categories as estimated using dynamic range boxes (R package "dynRB"). Heatmaps showing the portion of the hypervolume of habitat in y covered by the hypervolume of habitat in x. **A-C** show overlap of 6-dimensional hypervolumes aggregated as product (**A**), arithmetic (**B**) and geometric mean (**C**). **D-H** show overlap along individual dimensions. Note different scales for each heatmap. Habitat categories: palm (resting on palm leaves), grass (hanging from grass), branch (resting on branches and leaves), hang (hanging from branches and leaves) and litter (resting on the leaf litter, ground, logs and trunks).

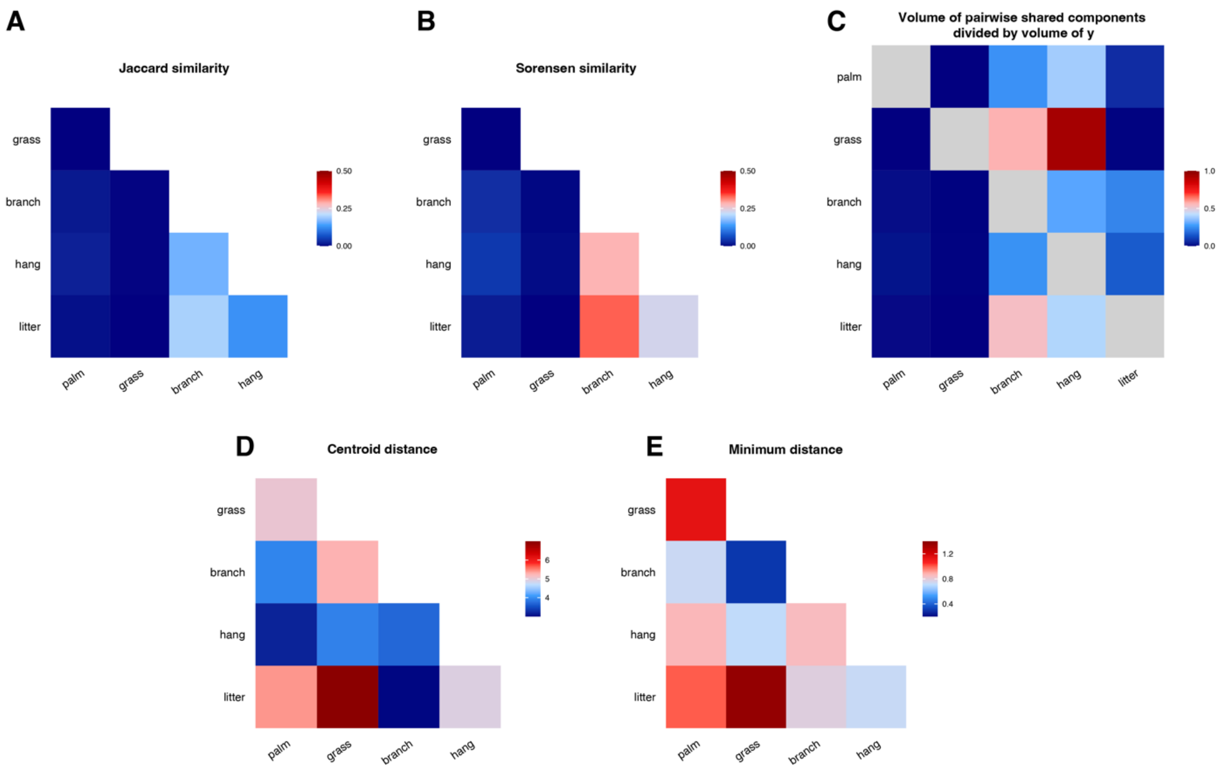


**Figure S10**: Hypervolume similarity (**A-C**) and distance (**D-E**) between different habitat categories as estimated using high-dimensional kernel density estimations (R package "hypervolume"). **A**: Pairwise Jaccard similarity quantifying the ratio of the intersection to the union of the hypervolumes. **B**: Pairwise Sørensen–Dice similarity index measuring the ratio between twice the size of the intersection between the hypervolumes and the sum of their individual sizes. **C**: Volume of shared components of the two hypervolumes divided by the size of the hypervolume of habitat in y. **D**: Pairwise Euclidean distance between the hypervolume centroids. **E**: Pairwise minimum Euclidean distance between two sets of random points comprising the two hypervolumes.

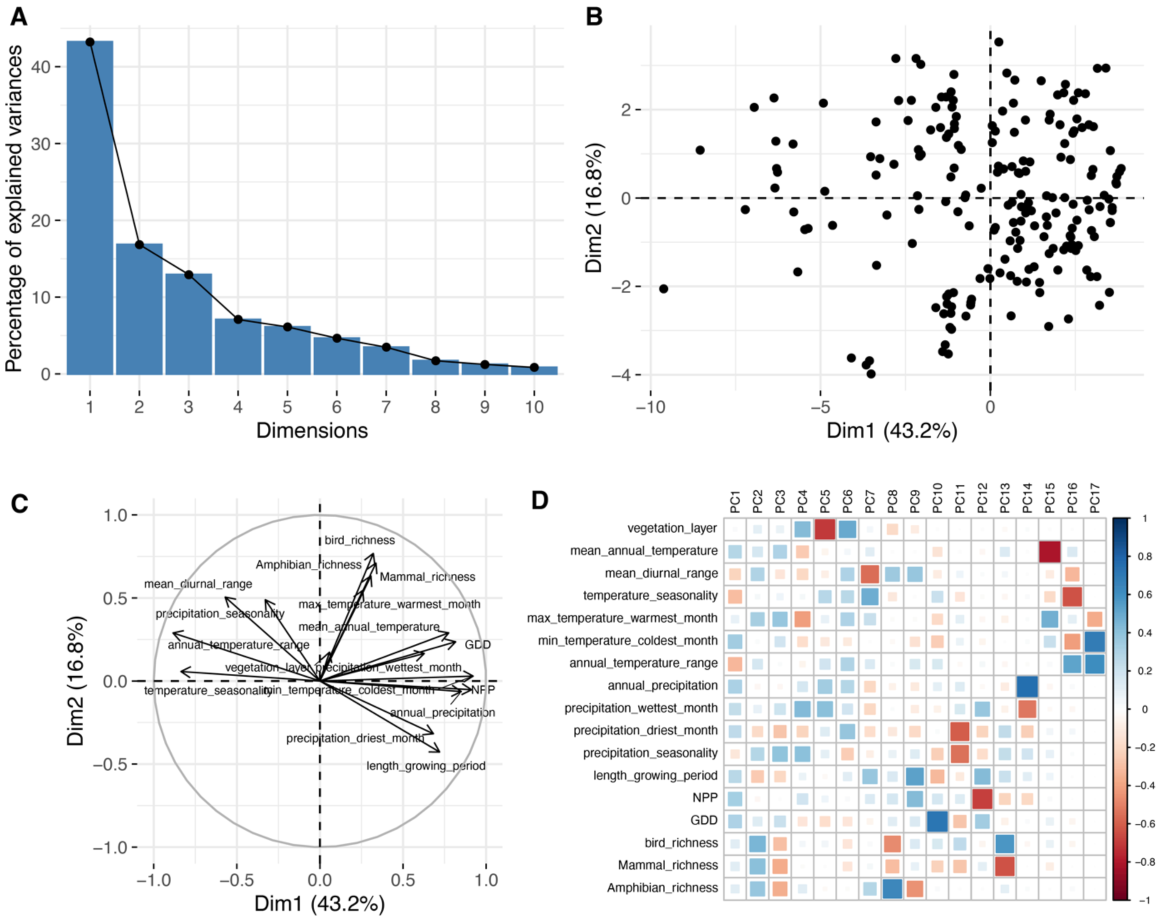


**Figure S11**: Results of the principal component analysis of 17 quantitative environmental variables. **A**: Proportion of total variance explained by each principal component. **B**: Position of each individual taxon along the first two principal components. **C**: Loading plot for the first principal components and the quantitative dataset features. **D**: correlation plot between the principal components and all the dataset features.

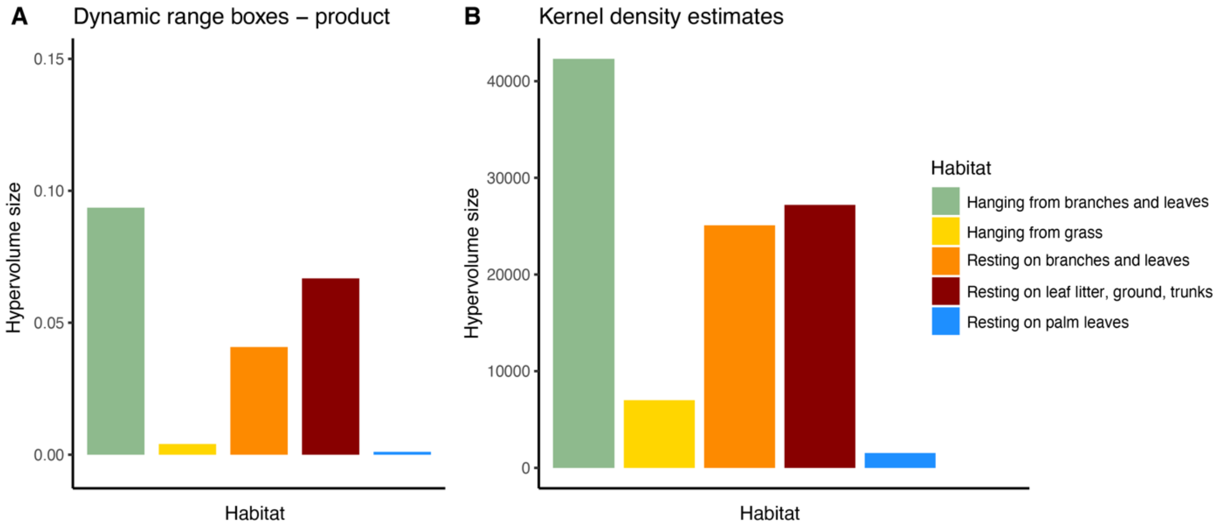


**Figure S12**: Habitat hypervolume size on the environmental space, estimated from dynamic range boxes and kernel density estimates using the first 6 axes (accounting for 91% of the total environmental variation).

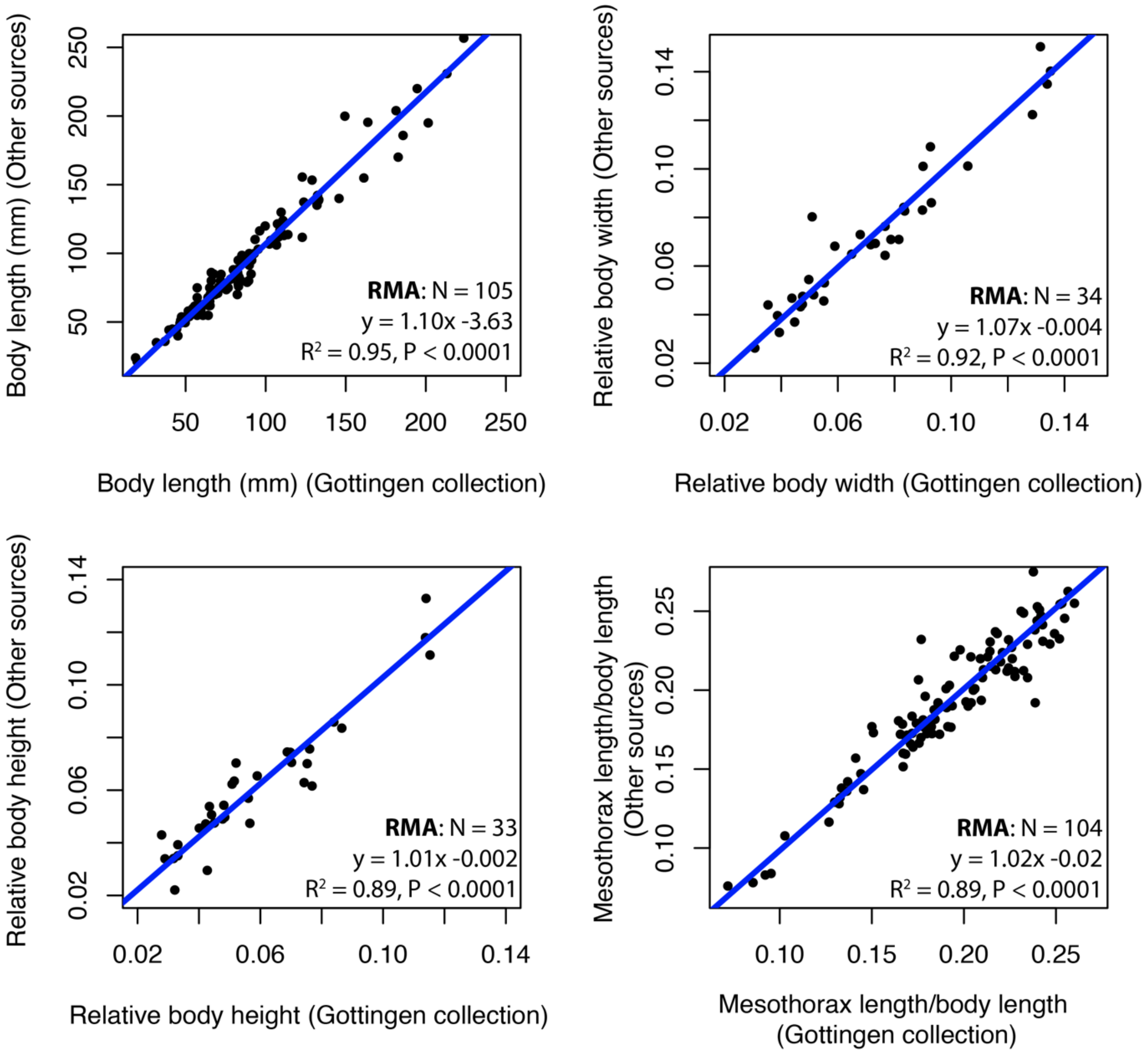


**Figure S13**: Comparisons between measurements obtained from our own collection at the University of Göttingen and measurements obtained from all other sources. Dots correspond to individual species for which both data types were collected. Results from reduced major axis regressions (RMA) are included in each corresponding panel.

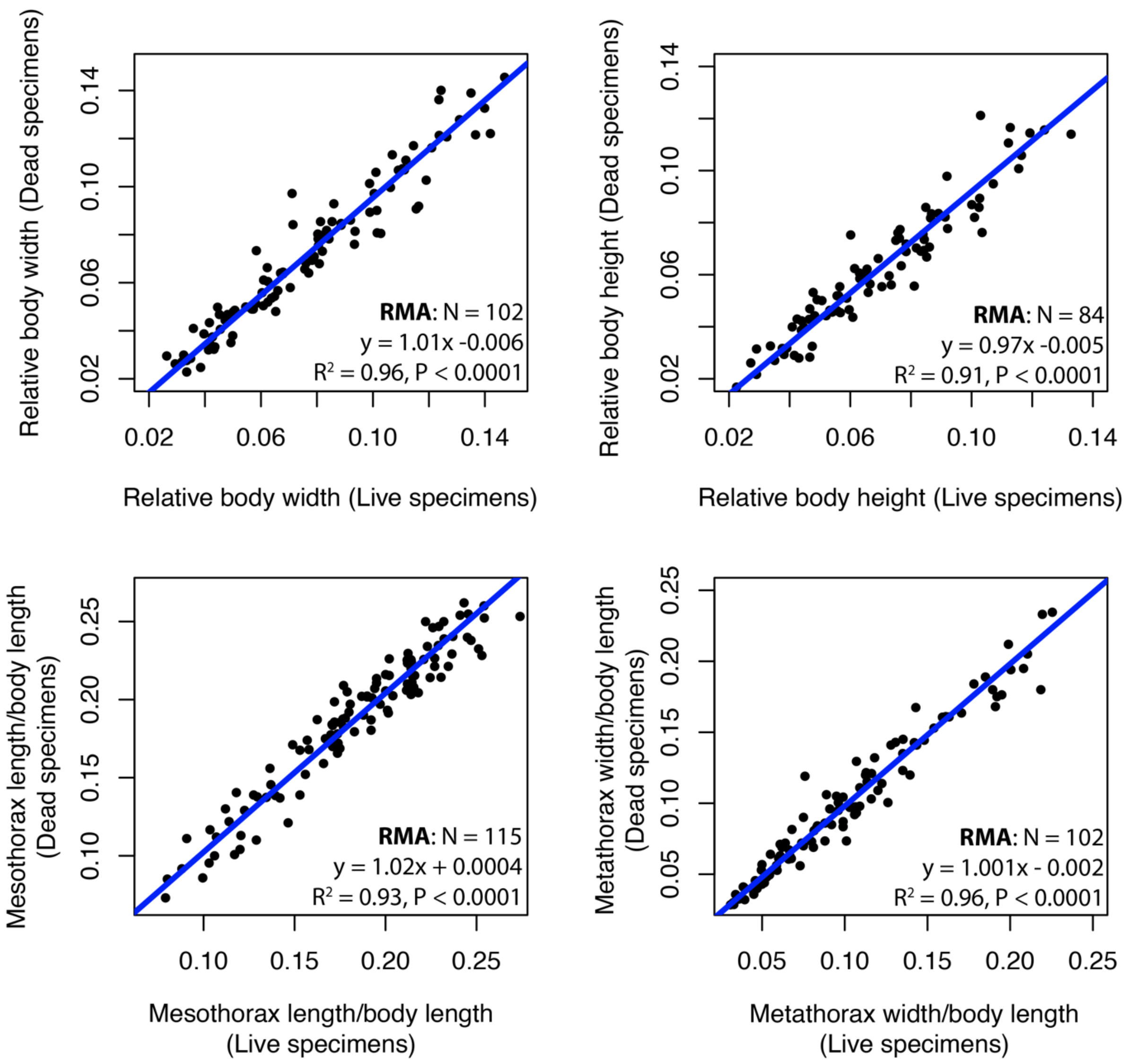


**Figure S14**: Comparisons between measurements obtained from pictures of dead specimens or live specimens. Dots correspond to individual species for which both data types were collected. Results from reduced major axis regressions (RMA) are included in each corresponding panel.

**Table S1:** Fossil calibrations used in the divergence time estimates. Fossils are numbered according to Figure 2.

| **Fossil** | **Life stage** | **Reference** | **Formation** | **Minimum age (Ma)** | **Calibration Node** | **Comments** |
|---|---|---|---|---|---|---|
| *Renphasma sinica* | Adult | (55) | Yixian formation, Liaonign, China | 122 | Root of tree, divergence of Embioptera and Phasmatodea (**fossil 1**) | Unambiguous crown group phasmatodean due to presence of the vomer between a pair of unsegmented cerci. |
| *Echinosomiscus primoticus* | Adult | (56) | Upper Cretaceous Amber, Myanmar | 98.8 | Divergence of Euphasmatodea and Timematodea (**fossil 2**) | Euphasmatodea with uncertain affinity (presumed male Neophasmatodea) |
| *Eophyllium messelensis* | Adult | (57) | Messel Germany deposits | 47 | Phyllinae (**fossil 3**) | Unambiguous crown leaf insect |
| *Eophasma spp.* | Eggs | (58) | Eocene Clarno Formation Nut Beds of Oregon | 44 | Pseudophasmatinae (**fossil 4**) | Euphasmatodean eggs most similar to Anisomorphini (Pseudophasmatinae) |
| *Clonistria sp.* | Eggs | (59) | Dominican amber, Hispaniola | 20-40 [40] | Diapheromerinae (**fossil 5**) | Unambiguous Diapheromerinae egg due to vesicular/matrix capitulum |
| *Malacomorpha sp.* | Eggs | (59) | Dominican amber, Hispaniola | 20-40 [20] | *Malacomorpha* (**fossil 6**) | Egg most similar to extant genus *Malacomorpha* (Pseudophasmatinae) |

**Table S2:** List of taxa represented in Figure 2.

1. *Timema californicum*
2. *Abrosoma festinatum*
3. *Dajaca napolovi*
4. *Dinophasma saginatum*
5. *Oreophoetes peruana*
6. *Diapheromera femorata*
7. *Cladomorphus phyllinus*
8. *Trychopeplus laciniatus*
9. *Cranidium gibbosum*
10. *Agathemera crassa*
11. *Metriophasma diocles*
12. *Paraprisopus antillarum*
13. *Anisomorpha buprestoides*
14. *Prisopus ariadne*
15. *Pseudophasma rufipes*
16. *Malacomorpha jamaicana*
17. *Peruphasma schultei*
18. *Epidares nolimentangere*
19. *Heteropteryx dilatata*
20. *Haaniella dehaanii*
21. *Aretaon asperrimus*
22. *Bacillus rossius*
23. *Spathomorpha adefa*
24. *Parectatosoma mocquerysi*
25. *Achrioptera punctipes*
26. *Cryptophyllium celebicum*
27. *Pulchriphyllium giganteum*
28. *Spinohirasea bengalensis*
29. *Kalocorinnis wegneri*
30. *Pseudodiacantha macklottii*
31. *Diesbachia tamyris*
32. *Phaenopharos struthioneus*
33. *Carausius morosus*
34. *Thaumatobactron guentheri*
35. *Eurycantha calcarata*
36. *Bactrododema hecticum*
37. *Taraxippus perezgelaberti*
38. *Diapherodes gigantea*
39. *Lamponius guerini*
40. *Phobaeticus serratipes*
41. *Parapachymorpha spiniger*
42. *Sceptrophasma hispidulum*
43. *Medaura sabriuscula*
44. *Clonaria conformans*
45. *Macrophasma biroi*
46. *Eurycnema goliath*
47. *Tropidoderus childrenii*
48. *Phasma gigas*

*49. Megacrania batesii*
*50. Extatosoma tiaratum*
*51. Dryococelus australis*
*52. Rhaphiderus spiniger*
*53. Apterograeffea reunionensis*
*54. Argosarchus horridus*
*55. Cnipsus rachis*

**Table S3:** List of taxa represented in Figure 4.

1. *Pulchriphyllium pulchrifolium*
2. *Nanophyllium frondosum*
3. *Microcanachus matileorum*
4. *Timema californicum*
5. *Prisopus ariadne*
6. *Ommatopseudes harmani*
7. *Epidares nolimetangere*
8. *Orestes bachmaensis*
9. *Agathemera crassa*
10. *Paraprisopus antillarum*
11. *Abrosoma festinatum*
12. *Dinophasma saginatum*
13. *Peruphasma schultei*
14. *Anisomorpha buprestoides (brown morph)*
15. *Macrophasma biroi*
16. *Diapherodes gigantea*
17. *Heteropteryx dilatata*
18. *Dryococelus australis*
19. *Eurycantha calcarata*
20. *Trychopeplus laciniatus*
21. *Extatosoma popa*
22. *Spinohirasea bengalensis*
23. *Cnipsus rachis*
24. *Taraxippus perezgelaberti*
25. *Parectatosoma mocquerysi*
26. *Aretaon asperrimus*
27. *Dimorphodes mancus*
28. *Orthomeria kangi*
29. *Oreophoetes peruana*
30. *Anisomorpha buprestoides (white morph)*
31. *Tropidoderus childrenii*
32. *Kalocorinnis wegneri*
33. *Anarchodes annulipes*
34. *Pseudophasma rufipes*
35. *Diesbachia tamyris*
36. *Spathomorpha adefa*
37. *Ctenomorpha marginipennis*
38. *Phobaeticus serratipes*
39. *Clonaria conformans*
40. *Clitarchus hookeri*
41. *Hyrtacus procerus*
42. *Phalces tuberculatus*
43. *Megacrania batesii*
44. *Graeffea leveri*
45. *Apterograeffea reunionensis*
46. *Phaenopharos struthioneus*
47. *Pharnacia ponderosa*
48. *Achrioptera spinosissima*

*49. Phasma gigas*
*50. Pseudodiacantha macklottii*
*51. Neopromachus wallacei*
*52. Pterinoxylus crassus*
*53. Argosarchus horridus*
*54. Monoiognosis spinosa*
*55. Sceptrophasma hispidulum*
*56. Clonopsis gallica*
*57. Carausius morosus*
*58. Hyrtacus tuberculatus*
59. *Diapheromera femorata*

**Table S4:** Description of the main distinctive morphological features of each ecomorph. The number of estimated independent origins on the phylogeny (see Fig 3C) and the name of the genera included are indicated. The main morphospace axes best describing the morphological "niche" of each ecomorph were identified using a random forest model predicting a given ecomorph versus all the others, given the first ten morphological axes (see Supplementary Materials and Methods). The importance of a given dimension was assessed by looking at the decrease in prediction accuracy after dropping each corresponding axes (as implemented in the function *randomForest*, R package "randomForest"). Only the main predictors are reported here.

| Ecomorph | # of Origins | Genera included | Main predicting morphospace axes |
|---|---|---|---|
| Wide leaf mimics | 1 | *Phyllium, Pulchriphyllium, Nanophyllium, Trolicaphyllium, Cryptophyllium, Chitoniscus* | PC2(43%): very flat body<br>PC1(17%): very wide body |
| Small crawlers | 2 | *Timema, Microcanachus* | PC3(26%): wingless<br>PC1(20%): relatively wide and stocky (also remarkably small) |
| Winged bark huggers | 3 | *Prisopus, Neoclides, Epicharmus* | PC6(22%): relatively small<br>PC3(20%): fully winged<br>PC1(14%): relatively wide |
| Diminutive spiny morphs | 3 | *Orestes, Epidares, Dares, Hoploclonia, Pterobrimus,, Labidiophasma, Ommatopseudes* | PC1 (23%): relatively wide and stocky.<br>PC2(17%): cylindrical body<br>PC4(14%): spiny and textured |
| Chinchemolles | 1 | *Agathemera* | PC6(27%): relatively massive and stocky insects<br>PC8(19%): dark<br>PC2(17%): cylindrical body |
| Apterous bark huggers | 7 | *Damasippoides, Pseudoleosthenes, Abrosoma, Dinophasma, Dajaca, Paraprisopus, Anisomorpha, Malacomorpha, Peruphasma, Pseudophasma, Thaumatobactron,* | PC1(16%): relatively wide and short legged<br>PC2(14%): cylindrical bodies<br>PC7(11%): dark |
| Lime green morphs | 6 | *Diapherodes, Venupherodes, Paramenexenus, Macrophasma, Monandroptera, Rhaphiderus, Cranidium.* | PC2(14%): relatively flattened<br>PC6(13%): relatively large<br>PC9(13%): long subgenital plate |
| Tree lobsters | 4 | *Eurycantha, Dryococelus, Canachus, Haaniella, Heteropteryx, Mearnsiana* | PC4 (25%): large and spiny<br>PC1 (22%): relatively wide |
| Spiny leaf mimics | 2 | *Extatosoma, Trychopeplus* | PC9 (26%): irregular body edges<br>PC4 (21%): spiny and textured |
| Spiny robust morphs | 11 | *Spinohirasea, Neohirasea, Oxyartes, Erinaceophasma, Neopromachus, Hyposcyrthus, Rhynchacris, Taraxippus, Lamponius, Dimorphodes, Micrarchus, Cnipsus, Creoxylus, Lobolibethra, Sungaya, Aretaon, Trachyaretaon, Brasidas, Parectatosoma* | PC1(27%): relatively wide<br>PC4(19%): spiny and/or textured |

| | | | |
|---|---|---|---|
| Aposematic morphs | 7 | *Oreophoetes, Tithonophasma, Anisomorpha, Pseudophasma, Orthomeria, Megacrania, Ophicrania* | PC8(42%): contrasting and conspicuous color pattern |
| Elongated leaf mimics | 3 | *Tropidoderus, Malandania, Parapodacanthus, Podacanthus* | PC3(45%): very large wings<br>PC5(12%): relatively small heads |
| Winged sticks | 2 | *Anarchodes, Diesbachia, Sipyloidea, Kalocorinnis, Trachythorax, Metriophasma, Malacomorpha, Pseudophasma* | PC3(32%): large wings<br>PC2(15%): cylindrical<br>PC6(15%): relatively small |
| Elongated sticks | 8 | *Phobaeticus, Phryganistria, Cuniculina, Staelonchodes, Lonchodes, Leprocaulinus, Baculofractum, Ctenomorpha, Bacteria, Spathomorpha* | PC1(46%): extremely long and elongated |
| Thin sticks | 10 | *Rhamphophasma, Clonaria, Ramulus, Medauroidea, Chondrostethus, Clitarchus, Hyrtacus,Orxines, Rhamphosipyloidea, Lopaphus,Leiophasma, Phalces,Phanocloidea* | PC1(31%): Very elongated<br>PC4(14%): smooth<br>PC3(12%): wingless |
| Large-headed sticks | 2 | *Megacrania, Ophicrania, Graeffea, Apterograeffea* | PC5(22%): relatively large head<br>PC10(20%): relatively elongated head |
| Large sticks | 5 | *Pharnacia, Tirachoidea, Phasmotaenia, Alienobostra, Cladomorphus, Phaenopharos* | PC1(16%): elongated<br>PC2(13%): tubular<br>PC6(11%): relatively large |
| Giant winged sticks | 4 | *Anchiale, Paronchestus, Acrophylla, Eurycnema, Phasma, Cigarophasma, Lopaphus, Bactrododema, Achrioptera* | PC3(30%): brachypterous<br>PC4(16%): spiny thorax<br>PC1(12%): elongated |
| Prickly sticks | 8 | *Argosarchus, Acanthoxyla, Asprenas, Mauritiophasma, Onchestus, Parapachymorpha, Agamemnon, Pterinoxylus, Neopromachus, Manduria, Pseudodiacantha, Centrophasma, Anisacantha, Antongilia* | PC4(18%): spiny and/or textured<br>PC8(17%): often brown<br>PC6(16%): often relatively small<br>PC1(12%): relatively elongated |
| Small sticks | 8 | *Ocnophiloidea, Diapheromera, Pseudosermyle, Caribbiopheromera, Bacillus, Clonopsis, Leptynia, Pijnackeria, Xylica, Zehntneria, Carausius, Hyrtacus, Lonchodes, Neopromachus, Parapachymorpha, Medaura, Sceptrophasma, Monoiognosis, Carlius.* | PC1(18%) : relatively elongated<br>PC5(14%): relatively small head<br>PC4(13%): smooth<br>PC2(11%): tubular |

**Table S5: Fits of multivariate models of character evolution.** Models were fit to the first 5 axes of the morphospace. Models with 2 regimes consider one habitat (indicated) as one regime, and the 4 remaining ones together as the other regime. The best fit model (bold) was determined by comparing AICc scores.

| | Model | LogLik | AICc | ΔAICc | Ak. weights |
|---|---|---|---|---|---|
| **1 regime** | BM | -1745.1 | 3531.0 | 238.4 | 0 |
| | EB | -1745.1 | 3533.1 | 240.5 | 0 |
| | OU1 | -1702.6 | 3477.8 | 185.2 | 0 |
| **2 regimes (habitats)** | BMm hanging from branches | -1715.0 | 3481.3 | 188.7 | 0 |
| | BMMm hanging from branches | -1660.5 | 3404.3 | 111.7 | 0 |
| | OUM hanging from branches | -1664.0 | 3411.4 | 118.8 | 0 |
| | BMm resting on ground/trunks | -1676.5 | 3404.2 | 11.6 | 0 |
| | BMMm resting on ground/trunks | -1633.3 | 3349.9 | 57.3 | 0 |
| | OUM resting on ground/trunks | -1662.4 | 3408.2 | 115.6 | 0 |
| | BMm resting on branches/leaves | -1691.1 | 3433.5 | 140.9 | 0 |
| | BMMm resting on branches/leaves | -1667.9 | 3419.1 | 126.5 | 0 |
| | OUM resting on branches/leaves | -1677.7 | 3438.8 | 146.1 | 0 |
| | BMm grass | -1740.3 | 3531.9 | 239.3 | 0 |
| | BMMm grass | -1713.4 | 3510.2 | 217.6 | 0 |
| | OUM grass | -1696.3 | 3475.9 | 183.2 | 0 |
| | BMm palm | -1736.6 | 3524.5 | 231.9 | 0 |
| | BMMm palm | -1717.4 | 3518.1 | 225.5 | 0 |
| | OUM palm | -1687.9 | 3459.2 | 166.6 | 0 |
| **5 regimes (habitats)** | BMm All | -1627.1 | 3337.6 | 45.0 | 0 |
| | **BMMm All** | **-1535.4** | **3292.6** | **0** | **1** |
| | OUM All | -1601.7 | 3319.7 | 27.1 | 0 |

For multiple-regime models, results are means from analyses using 100 stochastic character mapping reconstructions. BMm and BMMm allow the phylogenetic means to vary between different regimes (habitats). AICc, small-sample corrected Akaike Information Criterion; Ak., Akaike; BM, Brownian motion; EB, Early Burst; LogLik, log-likelihood; OU, Ornstein- Uhlenbeck.

**Table S6:** Fits of multivariate models of character evolution. Models were fit to the first 5 axes of the morphospace. Models with 20 regimes consider each ecomorph as one regime. The best fit model (with lowest AICc) is highlighted in bold.

| | Model | LogLik | AICc | ΔAICc | Ak. weights |
|---|---|---|---|---|---|
| **1 regime** | BM | -1745.1 | 3531.0 | 1139.9 | 0 |
| | EB | -1745.1 | 3533.1 | 1142.0 | 0 |
| | OU1 | -1702.6 | 3477.8 | 1086.6 | 0 |
| **20 regimes (ecomorphs)** | BMm All | -1181.5 | 2622.4 | 231.3 | 0 |
| | BMMm All | -909.7 | 3133.4 | 742.3 | 0 |
| | **OU** All | **-1046.5** | **2391.2** | **0** | **1** |

For multiple-regime models, results are means from analyses using 100 stochastic character mapping reconstructions. BMm and BMMm allow the phylogenetic means to vary between different regimes (habitats). AICc, small-sample corrected Akaike Information Criterion; Ak., Akaike; BM, Brownian motion; EB, Early Burst; LogLik, log-likelihood; OU, Ornstein- Uhlenbeck.

**Table S7: Morphological convergence metrics associated with independent habitat transitions.** For lineages including several taxa in our sample, pairwise metrics (C1-C4, Ct1-Ct4, θ) were calculated for all possible inter-group pairs of taxa and averaged. For these metrics, reported values represent means calculated across all pairs of independent transitions. P-values were calculated from 1000 Brownian motion simulations.

| | Resting on litter/trunks | Hanging from branches/leaves | Resting on branches/leaves | Hanging from grass | Resting on palm leaves |
|---|---|---|---|---|---|
| Number of independent origins | N = 15 | N = 4 | N = 14 | N = 4 | N = 2 |
| Wheatsheaf index *w* | **3.480 (p<0.001)** | 2.487 (p=0.20) | **2.665 (p=0.02)** | **6.236 (p<0.001)** | 2.843 (p=0.17) |
| C1 | **0.389 (p<0.001)** | 0.163 (p=0.27) | **0.153 (p=0.02)** | **0.583 (p<0.001)** | **0.268 (p=0.04)** |
| C2 | **2.23 (p<0.001)** | 0.885 (p=0.15) | **0.782 (p=0.02)** | **2.88 (p<0.001)** | **1.12 (p=0.02)** |
| C3 | **0.17 (p<0.001)** | 0.053 (p=0.48) | 0.065 (p=0.09) | **0.265 (p<0.001)** | **0.121 (p=0.05)** |
| C4 | **0.00004 (p<0.001)** | 0.00001 (p=0.64) | **0.0004 (p<0.001)** | 0.00002 (p=0.22) | 0.0002 (p=0.08) |
| Ct1 | **0.098 (p<0.001)** | -0.073 (p=0.40) | -0.227 (p=0.31) | -0.338 (p=0.35) | -0.022 (p=0.13) |
| Ct2 | **0.513 (p<0.001)** | -0.622 (p=0.81) | -0.840 (p=0.86) | **-0.176 (p=0.04)** | -0.057 (p=0.09) |
| Ct3 | **0.039 (p<0.001)** | -0.036 (p=0.53) | -0.071 (p=0.26) | **-0.043 (p=0.05)** | -0.007 (p=0.09) |
| Ct4 | **0.001 (p<0.001)** | -0.003 (p=0.81) | **-0.002 (p=0.02)** | -0.011 (p=0.42) | -0.001 (p=0.08) |
| Trajectory angle $\theta$ (°) | **40.50 (p<0.001)** | 84.26 (p=0.26) | **81.66 (p=0.01)** | **69.43 (p=0.05)** | **42.18 (p=0.007)** |

**Table S8: Effects of environmental distance and divergence time on morphological convergence.** Multiple matrix regressions (partial Mantel tests) testing the relative effect of ancestral state similarity, environmental distance and divergence time on morphological distance at present ($D_{tip}$), maximum morphological distance ($D_{max}$), C1 and trajectory angle (θ) between pairs of lineages that independently transitioned to resting on trunks or leaf litter (upper part) or to resting on branches and leaves (bottom part). Significant effects ($p<0.05$) are bolded.

| Dependent matrix | Independent matrices | Multiple $R^2$ | Coefficient | t | p |
|---|---|---|---|---|---|
| Transitions to resting on trunks or leaf litter | | | | | |
| $D_{tip}$ | Ancestral state | | 0.28 | 1.33 | 0.52 |
| | Environmental distance | 0.28 | 0.25 | 3.70 | **0.01** |
| | Divergence time | | 0.05 | 4.71 | **0.003** |
| $D_{max}$ | Ancestral state | | 0.39 | 1.77 | 0.45 |
| | Environmental distance | 0.27 | 0.07 | 1.01 | 0.55 |
| | Divergence time | | 0.06 | 5.45 | **0.002** |
| $C_1$ | Ancestral state | | -0.004 | -0.12 | 0.95 |
| | Environmental distance | 0.11 | -0.04 | -3.61 | **0.01** |
| | Divergence time | | -0.002 | -0.95 | 0.54 |
| Trajectory angle θ | Ancestral state | | -8.30 | -3.01 | 0.09 |
| | Environmental distance | 0.12 | 1.53 | 1.69 | 0.26 |
| | Divergence time | | 0.45 | 3.18 | **0.04** |
| Transitions to resting on branches and leaves | | | | | |
| $D_{tip}$ | Ancestral state | | -0.26 | -0.71 | 0.41 |
| | Environmental distance | 0.33 | 0.33 | 2.22 | **0.03** |
| | Divergence time | | 0.09 | 4.61 | **0.001** |
| $D_{max}$ | Ancestral state | | 0.10 | 0.41 | 0.63 |
| | Environmental distance | 0.30 | 0.13 | 1.33 | 0.23 |
| | Divergence time | | 0.06 | 4.34 | **0.002** |
| $C_1$ | Ancestral state | | 0.07 | 1.61 | 0.08 |
| | Environmental distance | 0.33 | -0.04 | -2.45 | **0.02** |
| | Divergence time | | -0.01 | -4.70 | **0.003** |
| Trajectory angle θ | Ancestral state | | -4.28 | -0.64 | 0.44 |
| | Environmental distance | 0.24 | 2.53 | 0.92 | 0.37 |
| | Divergence time | | 1.57 | 4.21 | **0.004** |

**Table S9:** List of taxa included in photographs in Figures S3-5.

| Picture # | Species | Origin | Source and photo credits |
|---|---|---|---|
| 1 | *Spathomorpha adefa* | Moramanga, Madagascar (wild) | CC-BY-NC-ND Davorka Kitonić & Josip Skejo |
| 2 | *Ctenomorpha marginipennis* | New South Wales, Australia (wild) | CC-BY-NC Paul Whitington |
| 3 | *Bacteria ploiaria* | Barro Colorado, Panama (wild) | CC-BY-NC John G. Phillips |
| 4 | *Phobaeticus serratipes* | Penang, Malaysia (wild) | CC-BY-NC Albert Kang |
| 5 | *Diapherodes martinicensis* | Martinique (captivity) | © Bruno Kneubühler, used by permission |
| 6 | *Paramenexenus laetus* | Tay Yen Tu NP, Vietnam (captivity) | © Bruno Kneubühler, used by permission |
| 7 | *Monandroptera acanthomera* | Réunion (wild) | © Nicolas Cliquennois, used by permission |
| 8 | *Cranidium gibbosum* | French Guiana (captivity) | CC-BY 3.0 Daniel Dittmar |
| 9 | *Orthomeria kangi* | Luzon, Philippines (wild) | CC-BY-NC Albert Kang |
| 10 | *Anisomorpha buprestoides* | Florida, USA (wild) | CC-BY-SA 3.0 Bugenstein |
| 11 | *Oreophoetes peruana* | Tarapoto, San Martín, Peru (wild) | © Romain Boisseau (author) |
| 12 | *Apterograeffea reunionensis* | Réunion (wild) | © Nicolas Cliquennois, used by permission |
| 13 | *Megacrania batesii* | Cape Tribulation, Queensland, Australia (wild) | © Romain Boisseau (author) |
| 14 | *Trachythorax maculliculis* | Thailand (captivity) | CC-BY-SA 3.0 Dragüs |
| 15 | *Pseudophasma phthisicum* | French Guiana (wild) | CC-BY-NC Manuel Ruedi |
| 16 | *Trychopeplus laciniatus* | Monteverde, Costa Rica (captivity) | © Bruno Kneubühler, used by permission |
| 17 | *Extatosoma tiaratum* | Queensland, Australia (captivity) | © Bruno Kneubühler, used by permission |
| 18 | *Diapheromera femorata* | Virginia, USA (wild) | CC-BY-2.0 Judy Gallagher |
| 19 | *Clonopsis gallica* | Jaén, Spain (wild) | CC BY 2.0 Ramón Portellano |
| 20 | *Dryococelus australis* | Ball's pyramid, Australia (captivity) | © Paul D. Brock, used by permission |
| 21 | *Haaniella dehaanii* | Sarawak, Borneo, Malaysia (captivity) | CC-BY-SA Drägüs |
| 22 | *Canachus alligator* | Mt Koghi, New Caledonia (captivity) | © Bruno Kneubühler, used by permission |
| 23 | *Eurycantha calcarata* | Kimbe, WNB, Papua New Guinea (wild) | © Romain Boisseau (author) |
| 24 | *Spinohirasea bengalensis* | Bach Ma, Vietnam (captivity) | © Bruno Kneubühler, used by permission |
| 25 | *Cnipsus rachis* | Mt Koghi, New Caledonia (wild) | CC-BY-NC Damien Brouste |
| 26 | *Parectatosoma sp.* | Moramanga, Madagascar (captivity) | © Bruno Kneubühler, used by permission |
| 27 | *Lamponius bocki* | Punta Cana, Dominican Republic (captivity) | © Bruno Kneubühler, used by permission |
| 28 | *Aretaon asperrimus* | Sabah, Borneo, Malaysia (wild) | CC-BY-NC Albert Kang |
| 29 | *Agathemera crassa* | Santiago, Chile (wild) | CC-BY-NC-SA Ariel Cabrera Foix |
| 30 | *Ommatopseudes harmani* | Tanah Rata, Peninsular Malaysia (captivity) | © Bruno Kneubühler, used by permission |
| 31 | *Epidares nolimetangere* | Sarawak, Borneo, Malaysia (captivity) | © Bruno Kneubühler, used by permission |
| 32 | *Labidiophasma rouxi* | Mt Humboldt, New Caledonia (wild) | CC-BY-NC Damien Brouste |
| 33 | *Timema boharti* | California, USA (wild) | CC-BY-NC John Christensen |
| 34 | *Microcanachus matileorum* | Yaté, New Caledonia (wild) | CC-BY-NC Damien Brouste |
| 35 | *Paraprisopus antillarum* | Guadeloupe (captivity) | © Bruno Kneubühler, used by permission |

| | | | |
|---|---|---|---|
| 36 | *Dinophasma saginatum* | Mulu NP, Sarawak, Borneo, Malaysia (captivity) | © Bruno Kneubühler, used by permission |
| 37 | *Pseudoleosthenes irregularis* | Ranomafana NP, Madagascar (wild) | © Paul Bertner, used by permission |
| 38 | *Anisomorpha buprestoides* | Florida, USA (wild) | CC-BY-NC Scott Ward |
| 39 | *Centrophasma hadrillum* | Bako NP, Sarawak, Borneo, Malaysia (captivity) | © Bruno Kneubühler, used by permission |
| 40 | *Argosarchus horridus* | Christchurch, New Zealand (wild) | CC-BY-NC Chris Morse |
| 41 | *Neopromachus sp.* | Popondetta, Oro Province, Papua New Guinea (wild) | © Romain Boisseau (author) |
| 42 | *Pulchriphyllium pulchrifolium* | Java, Indonesia (captivity) | © Romain Boisseau (author) |
| 43 | *Prisopus berosus* | Ocosingo, Chis, Mexico (wild) | CC-BY-NC Silvano LG |
| 44 | *Epicharmus marchali* | Mauritius (wild) | © Sylvain Hugel & Nicolas Cliquennois, used by permission |
| 45 | *Neoclides buescheri* | Bako NP, Sarawak, Borneo, Malaysia (captivity) | © Bruno Kneubühler, used by permission |
| 46 | *Tropidoderus childrenii* | New South Wales, Australia (wild) | CC-BY-NC-ND 2.0 David Midgley |
| 47 | *Malandania pulchra* | Queensland, Australia (wild) | CC-BY-NC Linda Rogan EntSocVic |
| 48 | *Phasmotaenia lanyuhensis* | Lanyuh Island (captivity) | © Bruno Kneubühler, used by permission |
| 49 | *Cladomorphus phyllinus* | Sao Paulo, Brasil (captivity) | © Bruno Kneubühler, used by permission |
| 50 | *Pharnacia ponderosa* | Mt Capotoan, Samar, Philippines (captivity) | © Bruno Kneubühler, used by permission |
| 51 | *Lopaphus sphalerus* | Qinnan, Qinzhou, Guangxi, China (captivity) | © Bruno Kneubühler, used by permission |
| 52 | *Achrioptera punctipes* | Madagascar (captivity) | © Bruno Kneubühler, used by permission |
| 53 | *Bactrododema hecticum* | Windhoek, Namibia (captivity) | © Bruno Kneubühler, used by permission |
| 54 | *Acrophylla wuelfingi* | Cairns, Queensland, Australia (wild) | CC-BY-NC Felix Fleck |

**Movie S1 (separate file).** Dynamic 2-D phylomorphospace. At each time step starting with the ancestral node of Phasmatodea, the position of each lineage on the morphospace was reconstructed and the morphotype of each lineage was predicted using a trained random forest model.

**Dataset S1 (separate file).** Genetic, morphological and ecological data used. A detailed description of the data is presented below.

| **Sheet** | **Column** | **Description** |
|---|---|---|
| Specimen_list_genbank<br><br>List of genetic sequences and accession numbers used to build phylogeny. | Subfamily Tribe | Subfamily name |
| | Genus | Genus name |
| | Species | Species name |
| | Taxon | Taxon name as appears in analyses |
| | Voucher | Specimen reference number |
| | Locality | Geographical origin of sample |
| | 12S, 16S, COI, COII, 18S, 28S, H3 | Genbank accession numbers for 12S, 16S, COI, COII, 18S, 28S, H3 sequences used to reconstruct phylogeny. |
| Morphodata_specimens<br><br>Morphological data of each specimen measured. See figure S1. | ID | Specimen ID number |
| | Within_species_ID | Specimen ID number for a given species |
| | Taxon | Taxon name |
| | Name_in_phylogeny | Taxon name as appears in phylogenetic tree |
| | Sex | Female (F) or Male (M) |
| | Image | Whether the measurements were obtained from photographs (y) or other sources (n) |
| | Scale_bar | Whether the photographs include a scale bar |
| | Image_orientation | Whether the photographs show the insect in dorsal, lateral or both views. |
| | source | Source of the photographs or data. May include the museum reference number, the name of the photographer and/or the scientific reference. |
| | Body_length_mm | Body length in mm |
| | Circ | Circularity of the insect's dorsal outline |
| | Solidity | Solidity of the insect's dorsal outline |
| | Av width | Average body width (relative to body length) |
| | Av height | Average body height (relative to body length) |
| | Av width without side extensions | Average body width excluding lateral abdominal extensions, such as those found in Phyllidae (relative to body length). |
| | Head length /body length | Head length relative to body length |
| | Head width /body length | Head width relative to body length |

| | | |
|---|---|---|
| | mesothorax front width /body length | Mesothorax width (front) relative to body length |
| | mesothorax rear width /body length | Mesothorax width (rear) relative to body length |
| | mesothorax length /body length | Mesothorax length relative to body length |
| | abdomen length (2 to 9 segment) middle /body length | Abdomen length (including the second to ninth segment) relative to body length |
| | abdomen width (2 segment front) /body length | Width of the 2nd abdominal segment relative to body length |
| | abdomen width (9 segment front) /body length | Width of the 9th abdominal segment relative to body length |
| | abdomen width (5 segment middle) /body length | Width of the 5th abdominal segment relative to body length |
| | metathorax width /body length | Metathorax width (middle) relative to body length |
| | metathorax length /body length | Metathorax length relative to body length |
| | mesothorax mid width /body length | Mesothorax width (middle) relative to body length |
| | femur length /body length | Front femur length relative to body length |
| | length_subgenital_plate /body length | Length of subgenital plate relative to body length |
| | wing_length /body_length | Length of hindwing relative to body length |
| | wing_area /body_length2 | Area of hindwing relative to body length (squared) |
| | green_morph | Whether the species displays a predominantly green morph (0 or 1) |
| | brown_morph | Whether the species displays a predominantly brown morph (0 or 1) |
| | other_color | Whether the species displays a predominantly non-green and non-brown morph (0 or 1) |
| | Textured_mesothorax | Whether the mesonotum is mostly smooth (no) or rough/spiny (yes) |
| | Textured_abdomen | Whether the abdomen is mostly smooth (no) or rough/spiny (yes) |
| | specimen_status | Whether the specimen was photographed/measured dead or alive |
| Morphospace_coordinates<br><br>Coordinates of each species in the morphospace (PC axes) | taxon | Taxon name as it appears in the phylogenetic tree |
| | dim1 – dim15 | Coordinates for each PC axis |
| | species | Taxon name |
| | clade | Major phylogenetic clade |
| | ecomorph | Ecomorph name |
| | vegetation_layer | Vegetation layer where species is typically found |
| | substrate | Typical resting substrate during the daytime |

| | | |
|---|---|---|
| | resting_posture | Typical resting posture during the daytime |
| | habitat_use | Combination of resting substrate and resting posture. |
| Habitat<br><br>Ecological data (species level) | species | Taxon name |
| | Current name | Latest taxon name |
| | ID_in_tree | Taxon name as it appears in the phylogenetic tree |
| | clade | Major phylogenetic clade |
| | oviposition_type | Female oviposition strategy |
| | vegetation_layer | Vegetation layer where species is typically found |
| | substrate | Typical resting substrate during the daytime |
| | resting_posture | Typical resting posture during the daytime |
| | habitat_use | Combination of resting substrate and resting posture. |
| | Reference_habitat | References for habitat classification |
| | ecomorph | Ecomorph name |
| | Country | Country of origin of the specimens sequenced |
| | Central_locality | Name of locality with most central latitude |
| | Central_latitude | Latitude of central locality |
| | Central_longitude | Longitude of central locality |
| | Lower_locality | Name of locality with lowest latitude |
| | Lower_latitude | Latitude of lower locality |
| | Lower_longitude | Longitude of lower locality |
| | Higher_locality | Name of locality with highest latitude |
| | Higher_latitude | Latitude of higher locality |
| | Higher_longitude | Longitude of higher locality |
| | References_locality | References for record locations |
| | mean_annual_temperature | Annual mean temperature at central locality (source: WorldClim) |
| | mean_diurnal_range | mean of monthly (maximum - minimum temperature) at central locality (source: WorldClim) |
| | temperature_seasonality | Temperature seasonality at central locality (standard deviation ×100) (source: WorldClim) |
| | max_temperature_warmest_month | Maximum Temperature of Warmest Month at central locality (source: WorldClim) |
| | min_temperature_coldest_month | Minimum Temperature of Coldest Month at central locality (source: WorldClim) |
| | annual_temperature_range | Temperature Annual Range (max_temperature_warmest_month – min_temperature_coldest_month) at central locality (source: WorldClim) |

| | | |
|---|---|---|
| | annual_precipitation | Annual Precipitation at central locality (source: WorldClim) |
| | precipitation_wettest_month | Precipitation of Wettest Month at central locality (source: WorldClim) |
| | precipitation_driest_month | Precipitation of Driest Month at central locality (source: WorldClim) |
| | precipitation_seasonality | Precipitation Seasonality (Coefficient of Variation) (source: WorldClim) |
| | length_growing_period | number of days during a year when temperatures are above 5°C and precipitation exceeds half the potential evapotranspiration at central locality (source: FAO map catalog) |
| | NPP | Net primary production of biomass (grams of dry matter per $m^2$ per year) at central locality (source: Center for Sustainability and the Global Environment, Nelson Institute for Environmental Studies at UW-Madison, USA) |
| | GDD | total annual growing degree days (heat units, based on a 5°C base temperature) at central locality (source: Center for Sustainability and the Global Environment, Nelson Institute for Environmental Studies at UW-Madison, USA) |
| | Bird_richness | Number of different bird species at central locality (source: iucnredlist.org) |
| | Mammal_richness | Number of different mammal species at central locality (source: iucnredlist.org) |
| | Amphibian_richness | Number of different amphibian species at central locality (source: iucnredlist.org) |
| | elevation_central_location | Elevation (m) at central locality |

**SI References**


1. R Core Team, R: A Language and Environment for Statistical Computing (2023).
2. B. Misof, *et al.*, Phylogenomics resolves the timing and pattern of insect evolution. *Science (1979)* **346**, 763–767 (2014).
3. J. A. Robertson, S. Bradler, M. F. Whiting, Evolution of oviposition techniques in stick and leaf insects (Phasmatodea). *Front Ecol Evol* **6**, 1–15 (2018).
4. S. Bank, *et al.*, Reconstructing the nonadaptive radiation of an ancient lineage of ground-dwelling stick insects (Phasmatodea: Heteropterygidae). *Syst Entomol* **46**, 487–507 (2021).

5. S. Bradler, J. A. Robertson, M. F. Whiting, A molecular phylogeny of Phasmatodea with emphasis on Necrosciinae, the most species-rich subfamily of stick insects. *Syst Entomol* **39**, 205–222 (2014).
6. F. Glaw, O. Hawlitschek, A. Dunz, J. Goldberg, S. Bradler, When giant stick insects play with colors: Molecular phylogeny of the Achriopterini and description of two new splendid species (Phasmatodea: Achrioptera) from Madagascar. *Front Ecol Evol* **7** (2019).
7. S. Bradler, N. Cliquennois, T. R. Buckley, Single origin of the Mascarene stick insects: ancient radiation on sunken islands? *BMC Evol Biol* **15**, 1–10 (2015).
8. G. Forni, *et al.*, Macroevolutionary Analyses Provide New Evidences of Phasmids Wings Evolution as a Reversible Process. *bioRxiv* (2021) https:/doi.org/10.1101/2020.10.14.336354.
9. E. S. Wright, Using DECIPHER v2. 0 to analyze big biological sequence data in R. *R J* **8** (2016).
10. E. Paradis, K. Schliep, ape 5.0: an environment for modern phylogenetics and evolutionary analyses in R. *Bioinformatics* **35**, 526–528 (2019).
11. V. Ranwez, E. J. P. Douzery, C. Cambon, N. Chantret, F. Delsuc, MACSE v2: Toolkit for the Alignment of Coding Sequences Accounting for Frameshifts and Stop Codons. *Mol Biol Evol* **35**, 2582–2584 (2018).
12. K. Schliep, T. Jombart, Z. Kamvar Namir, E. Archer, R. Harris, apex: Phylogenetic Methods for Multiple Gene Data (2020).
13. R. Lanfear, P. B. Frandsen, A. M. Wright, T. Senfeld, B. Calcott, PartitionFinder 2: New Methods for Selecting Partitioned Models of Evolution for Molecular and Morphological Phylogenetic Analyses. *Mol Biol Evol* **34**, 772–773 (2017).
14. M. A. Miller, W. Pfeiffer, T. Schwartz, The CIPRES Science Gateway: A Community Resource for Phylogenetic Analyses in *Proceedings of the 2011 TeraGrid Conference on Extreme Digital Discovery - TG '11*, (ACM Press, 2011) https:/doi.org/10.1145/2016741.
15. R. Bouckaert, *et al.*, BEAST 2: A Software Platform for Bayesian Evolutionary Analysis. *PLoS Comput Biol* **10**, e1003537 (2014).
16. R. R. Bouckaert, A. J. Drummond, bModelTest: Bayesian phylogenetic site model averaging and model comparison. *BMC Evol Biol* **17**, 1–11 (2017).
17. E. Tihelka, C. Cai, M. Giacomelli, D. Pisani, P. C. J. Donoghue, Integrated phylogenomic and fossil evidence of stick and leaf insects (Phasmatodea) reveal a Permian–Triassic co-origination with insectivores. *R Soc Open Sci* **7**, 201689 (2020).
18. A. Rambaut, A. J. Drummond, D. Xie, G. Baele, M. A. Suchard, Posterior Summarization in Bayesian Phylogenetics Using Tracer 1.7. *Syst Biol* **67**, 901 (2018).
19. F. Seow-Choen, *A Taxonomic Guide to the Stick Insects of Borneo: Including New Genera and Species*, Natural Hi (2016).
20. F. Seow-Choen, *A Taxonomic Guide to the Stick Insects of Singapore* (Natural History Publications (Borneo), 2017).
21. F. Seow-Choen, *Phasmids of Peninsular Malaysia and Singapore* (Natural History Publications (Borneo), 2005).
22. F. Seow-Choen, *A Taxonomic Guide to the Stick Insects of Sumatra, Volume 1* (Natural History Publications (Borneo), 2018).
23. P. D. Brock, J. W. Hasenpusch, *The complete field guide to stick and leaf insects of Australia* (CSIRO publishing, 2009).
24. P. D. Brock, T. H. Büscher, E. Baker, Phasmida Species File Online. *Version 5.0/5.0* (2021).
25. C. A. Schneider, W. S. Rasband, K. W. Eliceiri, NIH Image to ImageJ: 25 years of image analysis. *Nat Methods* **9**, 671–675 (2012).
26. M. Chavent, V. Kuentz-Simonet, A. Labenne, J. Saracco, Multivariate Analysis of Mixed Data: The R Package PCAmixdata. *ArXiv* (2017).
27. J. C. Carvalho, P. Cardoso, Decomposing the Causes for Niche Differentiation Between Species Using Hypervolumes. *Front Ecol Evol* **8** (2020).
28. L. J. Revell, Size-correction and principal components for interspecific comparative studies. *Evolution (N Y)* **63**, 3258–3268 (2009).

29. L. J. Revell, phytools: An R package for phylogenetic comparative biology (and other things). *Methods Ecol Evol* **3**, 217–223 (2012).
30. S. Bradler, T. R. Buckley, “Biodiversity of Phasmatodea” in *Insect Biodiversity: Science and Society*, R. G. Foottit, P. H. Adler, Eds. (Wiley-Blackwell, 2018), pp. 281–313.
31. G. O. Bedford, Biology and ecology of the Phasmatodea. *Annu Rev Entomol* **23**, 125–149 (1978).
32. P. D. Brock, T. H. Büscher, *Stick and Leaf-Insects of the World* (NAP Editions, 2022).
33. S. Datta, S. Datta, Methods for evaluating clustering algorithms for gene expression data using a reference set of functional classes. *BMC Bioinformatics* **7**, 1–9 (2006).
34. G. Brock, V. Pihur, S. Datta, S. Datta, clValid: An R Package for Cluster Validation. *J Stat Softw* **25**, 1–22 (2008).
35. A. Liaw, M. Wiener, Classification and Regression by randomForest. *R News* **2**, 18–22 (2002).
36. A. L. Pigot, *et al.*, Macroevolutionary convergence connects morphological form to ecological function in birds. *Nat Ecol Evol* **4**, 230–239 (2020).
37. R. R. Junker, J. Kuppler, A. C. Bathke, M. L. Schreyer, W. Trutschnig, Dynamic range boxes – a robust nonparametric approach to quantify size and overlap of n-dimensional hypervolumes. *Wiley Online Library* **7**, 1503–1513 (2016).
38. B. Blonder, *et al.*, hypervolume: high dimensional geometry, set operations, projection, and inference using kernel density estimation, support vector machines, and convex hulls (2023).
39. A. Liaw, M. Wiener, Classification and regression by randomForest. *R News* **2**, 18–22 (2002).
40. K. K. Nordén, *et al.*, Melanosome diversity and convergence in the evolution of iridescent avian feathers—Implications for paleocolor reconstruction. *Evolution (N Y)* **73**, 15–27 (2019).
41. J. Clavel, G. Escarguel, G. Merceron, mvmorph: an r package for fitting multivariate evolutionary models to morphometric data. *Methods Ecol Evol* **6**, 1311–1319 (2015).
42. K. Arbuckle, C. M. Bennett, M. P. Speed, A simple measure of the strength of convergent evolution. *Methods Ecol Evol* **5**, 685–693 (2014).
43. K. Arbuckle, A. Minter, Windex: Analyzing convergent evolution using the wheatsheaf index in R. *Evolutionary Bioinformatics* **2015**, 11–14 (2015).
44. C. T. Stayton, The definition, recognition, and interpretation of convergent evolution, and two new measures for quantifying and assessing the significance of convergence. *Evolution (N Y)* **69**, 2140–2153 (2015).
45. D. M. Grossnickle, *et al.*, Challenges and advances in methods for measuring phenotypic convergence. *bioRxiv*, 2022.10.18.512739 (2023).
46. S. Schlager, *Morpho and Rvcg - Shape Analysis in R*, G. Zheng, S. Li, G. Szekely, Eds. (Academic Press, 2017) https:/doi.org/10.1016/B978-0-12-810493-4.00011-0.
47. S. E. Fick, R. J. Hijmans, WorldClim 2: new 1-km spatial resolution climate surfaces for global land areas. *International Journal of Climatology* **37**, 4302–4315 (2017).
48. H. van Velthuizen, *et al.*, *Mapping biophysical factors that influence agricultural production and rural vulnerability* (Food and Agriculture Organization of the United Nations and International Institute for Applied Systems Analysis, 2007).
49. J. B. Losos, Convergence, adaptation, and constraint. *Evolution (N Y)* **65**, 1827–1840 (2011).
50. D. L. Mahler, M. G. Weber, C. E. Wagner, T. Ingram, Pattern and Process in the Comparative Study of Convergent Evolution. *Am Nat* **190**, S13–S28 (2017).
51. D. M. Grossnickle, *et al.*, Incomplete convergence of gliding mammal skeletons. *Evolution (N Y)* **74**, 2662–2680 (2020).
52. D. C. Collar, J. S. Reece, M. E. Alfaro, P. C. Wainwright, R. S. Mehta, Imperfect morphological convergence: Variable changes in cranial structures underlie transitions to durophagy in moray eels. *American Naturalist* **183** (2014).
53. S. Castiglione, *et al.*, A new, fast method to search for morphological convergence with shape data. *PLoS One* **14**, e0226949 (2019).

54. M. P. Speed, K. Arbuckle, Quantification provides a conceptual basis for convergent evolution. *Biological Reviews* **92**, 815–829 (2017).
55. A. Nel, E. Delfosse, A new Chinese Mesozoic stick insect. *Acta Palaeontol Pol* **56**, 429–432 (2011).
56. M. S. Engel, B. Wang, A. S. Alqarni, A thorny, "anareolate" stick-insect (Phasmatidae s.l.) in Upper Cretaceous amber from Myanmar, with remarks on diversification times among Phasmatodea. *Cretac Res* **63**, 45–53 (2016).
57. S. Wedmann, S. Bradler, J. Rust, The first fossil leaf insect: 47 Million years of specialized cryptic morphology and behavior. *Proc Natl Acad Sci U S A* **104**, 565–569 (2007).
58. J. T. C. Sellick, Phasmida (stick insect) eggs from the Eocene of Oregon. *Palaeontology* **37**, 913–921 (1994).
59. G. Poinar, A walking stick, *Clonistria dominicana* n. sp. (phasmatodea: Diapheromeridae) in Dominican amber. *Hist Biol* **23**, 223–226 (2011).
