## Supplementary figures and images for "Divergence time and environmental similarity predict the strength of morphological convergence in stick and leaf insects"

### Movie S1

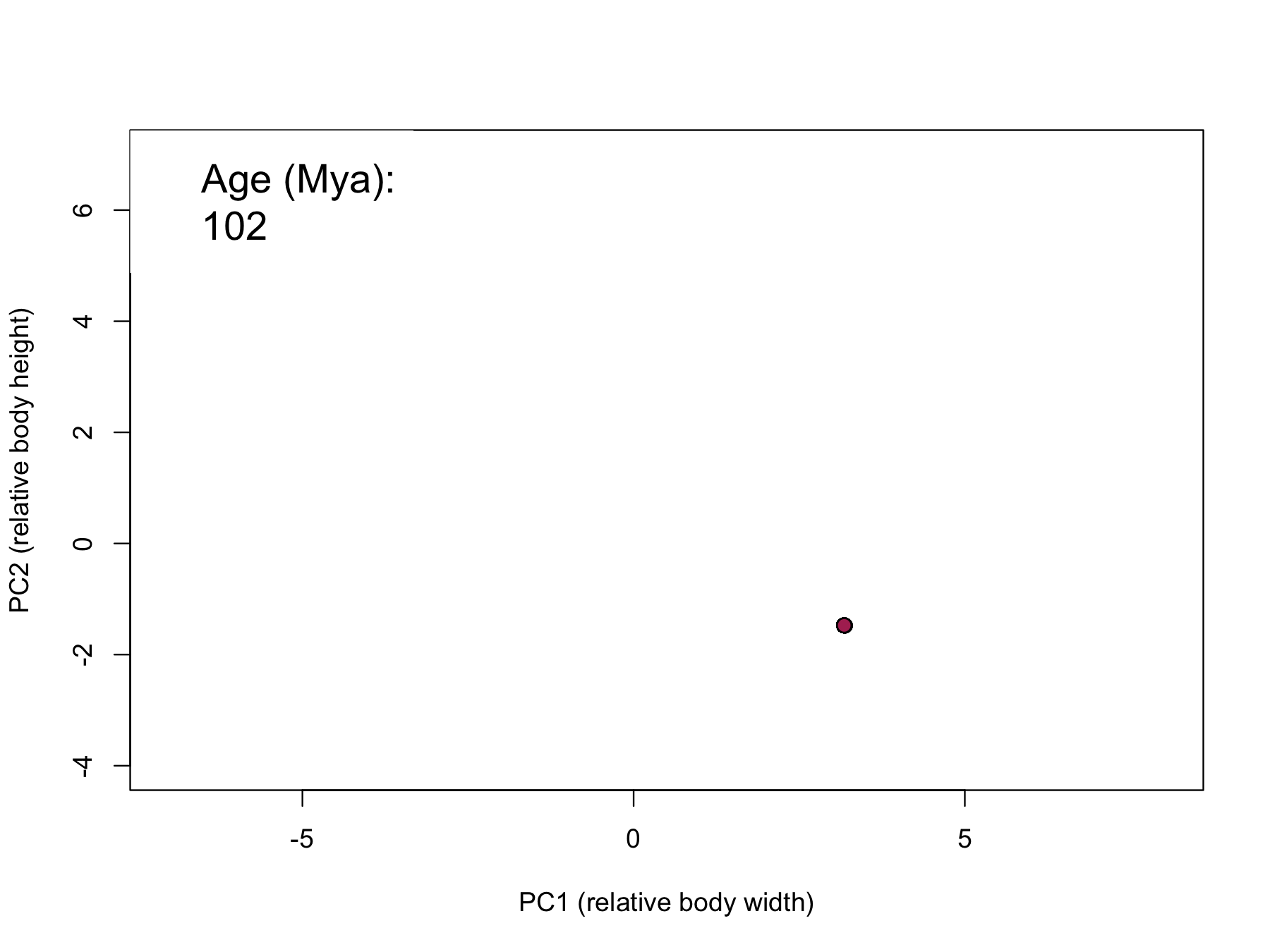
